# Doublecortin mutation disrupts lattice-dependent proximity to reveal cytoskeletal networks in developing neurons

**DOI:** 10.64898/2026.09.11.750980

**Authors:** Brandi L. Kelly, Justin A.G. Haineault, Lisa M. Munter, Gary J. Brouhard, Jean-François Trempe

**Affiliations:** Department of Biochemistry, McGill University, Montréal, QC, Canada; Centre de Recherche en Biologie Structurale, McGill University, Montréal, QC, Canada; Department of Pharmacology and Therapeutics, McGill University, Montréal, QC, Canada; Department of Biology, McGill University, Montréal, QC, Canada; Brain Repair and Integrative Neuroscience (BRaIN) Program, Research Institute of the McGill University Health Center, Montréal, QC, Canada

**Keywords:** Doublecortin, BioID, proteomics, iPSC-derived neurons, neuronal differentiation, microtubule-associated proteins, cytoskeletal proteins

## Abstract

Doublecortin (DCX) is a microtubule-associated protein (MAP) required for neuronal migration and cortical development. However, DCX is not sufficient to direct these processes on its own. Instead, nucleokinesis, growth cone advancement, collateral branching, and polarity depend on the coordination of different MAPs and cytoskeletal proteins, including DCX, to define distinct polymer properties and cytoskeletal interfaces. While many of these proteins have been investigated for their mechanistic cytoskeletal engagement, the spatiotemporal proteomic landscape of developing neurons remains largely uncharacterized. Here, we explore DCX-centric cytoskeletal networks using BioID accumulated throughout the differentiation of cortical neurons from induced pluripotent stem cells (iPSCs). Using the morphological staging of these cells as temporal landmarks, we generate a time-resolved proteomic atlas that charts DCX’s molecular environment throughout differentiation. From this atlas, we identify a cluster of cytoskeletal proteins whose persistent proximity to DCX is disrupted with the disease-associated mutation, DCX p.Arg178Leu (R178L). Consistent with R178L’s diminished cooperativity, we find many of these species are implicated in modulating microtubule stability and intracellular transport, including proteins EML4 and MAP7D1. Together, our findings place DCX at the interface of distinct microtubule populations and cytoskeletal systems throughout neuronal development.

## Introduction

Doublecortin (DCX) is a microtubule-associated protein (MAP) that regulates the migration and morphology of developing neurons (Des Portes, V. *et al*., 1998a; Gleeson, J. G. *et al*., 1998; Gleeson, J. G. *et al*., 1999a). DCX was first identified as a causative gene in patients with lissencephaly and subcortical band heterotopia (Des Portes, V. *et al*., 1998a; Gleeson, J. G. *et al*., 1998). Since then, lissencephaly-associated mutations have been extensively characterized: in cellular systems, for their effects on neuronal morphology and microtubule organization (Gleeson, J. G. *et al*., 1999b; Sébastien, M. *et al*., 2025; Yap, C. C. *et al*., 2016), and in purified reconstitution assays, for their effects on microtubule-binding kinetics (Bechstedt, S. & Brouhard, Gary J., 2012). However, the precise function of DCX in regulating neuronal development remains unclear.

Beyond its role as a microtubule-binding protein, DCX is implicated in several cytoskeletal activities, including polarity maintenance (Koizumi, H. *et al*., 2006; Sébastien, M. *et al*., 2025), the bundling of actin and microtubules (Toriyama, M. *et al*., 2012; Tsukada, M. *et al*., 2005), positioning of the nucleus and Golgi (Koizumi, H. *et al*., 2006; Li, P. *et al*., 2021; Stouffer, M. A. *et al*., 2022), regulation of axonal branching (Kappeler, C. *et al*., 2006; Slepak, T. I. *et al*., 2012; Tint, I. *et al*., 2009), and growth cone advancement in neurons (Dema, A. *et al*., 2024; Tint, I. *et al*., 2009). Moreover, its localization and participation in these activities is routinely modulated by phosphorylation (Bielas, S. L. *et al*., 2007; Bott, C. J. *et al*., 2020; Jin, J. *et al*., 2010). Even so, few proteins are reported to interact with DCX in ways that explain these diverse cellular functions. Among the best characterized interactions, most instead support a role for DCX in regulating intracellular transport (Fu, X. *et al*., 2022; Li, P. *et al*., 2021; Liu, Judy S. *et al*., 2012; Monroy, B. Y. *et al*., 2018).

A challenge in reconciling DCX’s diverse cellular and biophysical activities can be attributed to its structure. Although both tandem globular "DC" domains bind microtubules, they exhibit distinct structural and biochemical properties. DC1 adopts a stable ubiquitin-like fold with a preference for compacted (GDP-bound) lattice conformations, whereas DC2 behaves as a molten globule with increased affinity for expanded (GTP-bound) lattice states (Kim, M. H. *et al*., 2003; Manka, S. W. & Moores, C. A., 2020; Uversky, V. N., 2002). These structured domains are also separated and flanked by three intrinsically disordered regions (Cierpicki, T. *et al*., 2006), features that have historically complicated the structural characterization of DCX (Burger, D. *et al*., 2016; Kim, M. H. *et al*., 2003; Rufer, A. C. *et al*., 2018). More recently, integrative structural studies have shown that these disordered elements may allow DC2 to adopt conformational states that depend on DC1-microtubule engagement (Rafiei, A. *et al*., 2022). Together with its potential for lattice-dependent DC2 self-association (Rafiei, Cruz Tetlalmatzi et al. 2022), these observations suggest that reported DCX functions may be dependent on local composition of the cytoskeleton.

Consistent with this hypothesis, purified reconstitution assays demonstrate that DCX stabilizes microtubules, binding at the interface between four tubulin dimers (Bechstedt, S. *et al*., 2012; Manka, S. W. *et al*., 2020; Moores, C. A. *et al*., 2004). Bulk microtubule stabilization is not readily observed in neurons, where DCX functions within dense networks of other MAPs that promote remodeling (Arpag, G. *et al*., 2020; Bieling, P. *et al*., 2007; McAlear, T. S. & Bechstedt, S., 2022). Instead, recent work suggests that DCX modulates the responsiveness of remodeled microtubules to local actin remodeling (Dema, A. *et al*., 2024), echoing earlier evidence for functional crosstalk between DCX and the actin cytoskeleton (Fu, X. *et al*., 2013). We therefore hypothesized that these diverse behaviors are determined through developmental programming and cytoskeletal proximity, which could be probed through DCX-based proximity labeling.

DCX is specifically expressed in immature neuronal populations throughout development (Gleeson, J. G. *et al*., 1999a). Thus, to probe DCX’s native spatiotemporal environment, we used CRISPR-Cas9-engineered induced pluripotent stem cell (iPSC) lines to perform BioID proximity labeling throughout the differentiation of neurogenin-2 (NGN2)-derived cortical neurons (Antonicka, H. *et al*., 2020; Branon, T. C. *et al*., 2018; Hulme, A. J. *et al*., 2022; Muñoz-Estrada, J. *et al*., 2026). Using an accumulated-labeling approach coupled with quantitative proteomics, we generated a cytoskeletal “proximity” atlas charting the relative intensity of nearby proteins at different points of differentiation. In addition to the broad cytoskeletal remodeling that occurs between migration- and polarization-like stages, a cluster of cytoskeletal proteins and MAPs exhibited reduced proximity to p.Arg178Leu (R178L), a DCX mutation which causes lissencephaly (Bechstedt, S. *et al*., 2012; Des Portes, V. *et al*., 1998b; Gleeson, J. G. *et al*., 1999b). Among those with R178L sensitivity, we found that EML4 and MAP7D1 exhibited a persistent proximity to DCX throughout differentiation. However, while both proteins exhibit compartmental enrichment patterns that overlap with DCX, these patterns are themselves divergent. Together, these results suggest that while EML4 and MAP7D1 are in proximity to DCX, they do not necessarily occupy the same microtubule networks. Our approach therefore resolves the DCX-proximal proteome along two complementary dimensions: R178L mutation identifies cytoskeletal neighborhoods that are sensitive to altered microtubule binding, while spatial analysis distinguishes functionally overlapping networks that remain compartmentally distinct.

## Results

### Cell line engineering and validation of morphological staging in NGN2-derived cortical neurons

To elucidate DCX’s native cytoskeletal environment throughout cortical differentiation, we engineered an iPSC line to express DCX fused with a hemagglutinin (HA)-tagged miniTurboID enzyme (**Figure 1 supplement 1A, 1B**). The parental iPSC line contained a doxycycline-inducible NGN2 promoter for directed differentiation of progenitors towards a cortical fate, within 96 hours, or 4 days *in vitro* (DIV; **Figure 1A**, **Figure 1 supplement 1C**) (Hulme, A. J. *et al*., 2022; Muñoz-Estrada, J. *et al*., 2026). Consistent with the morphological progression of alternative models, the onset of DCX expression preceded bipolar axis formation, beginning at DIV2, roughly marking neural progenitor stage that is bypassed with NGN2-induction (Hindley, C. *et al*., 2012; Muñoz-Estrada, J. *et al*., 2026; Sébastien, M. *et al*., 2025). Likewise, DCX expression increased throughout differentiation alongside B3-tubulin (TUBB3), a marker of neuronal development (Sébastien, M. *et al*., 2025). While expression of the ∼75 kDa fusion protein (DCX-mTID) was reduced compared to the unedited control (CTRL0), TUBB3 expression was consistent across cell lines, onset of DCX expression remained the same, and anti-biotin staining of lysates confirmed enzymatic activity of the fused biotin ligase (**Figure 1B**).

**Figure 1.**
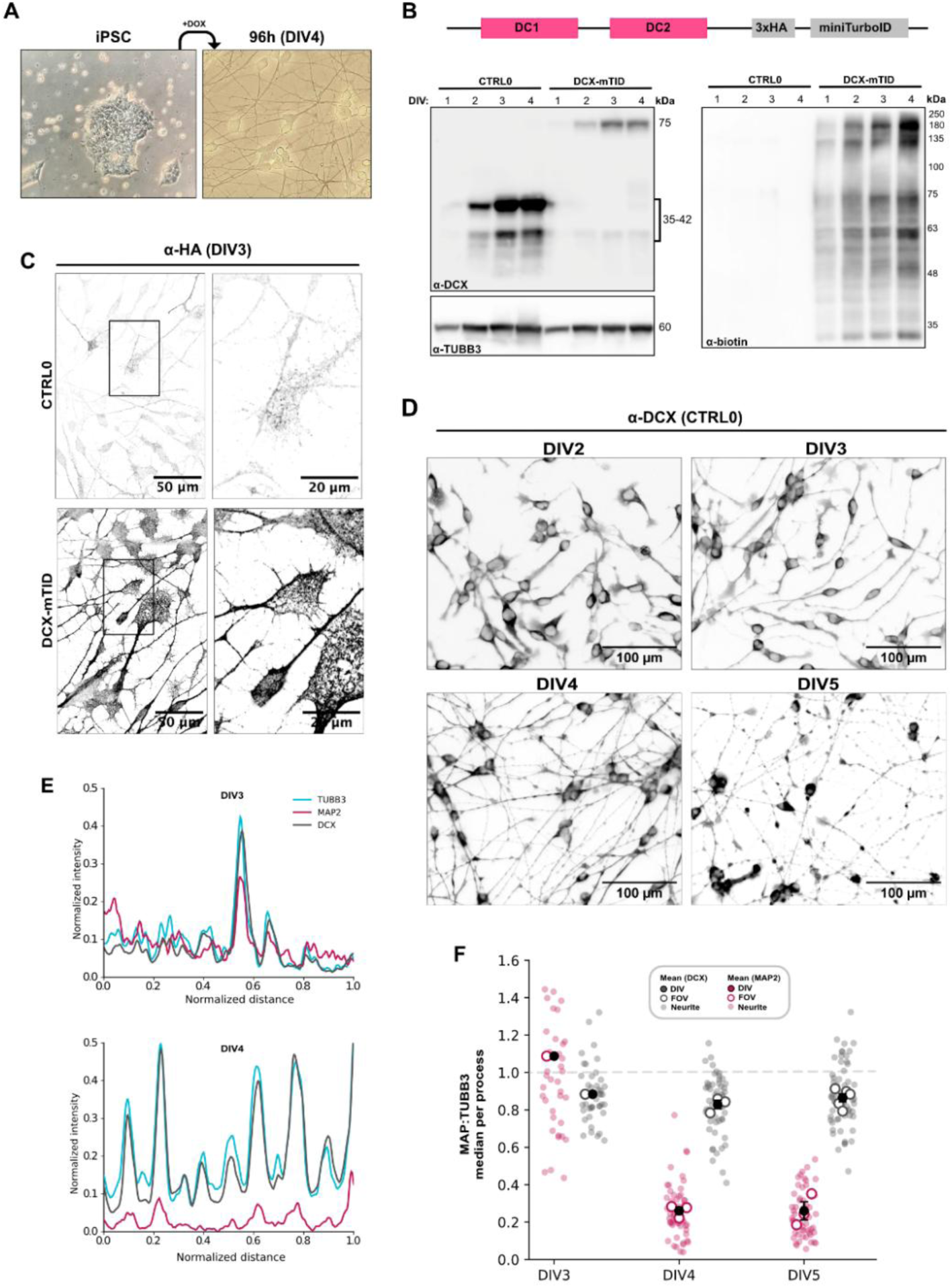
(with 2 supplements) – *Cell line engineering and validation of morphological staging in NGN2-derived cortical neurons*. **(A)** Schematic illustrates morphology of Bi0Ni010-C13 iPSCs before and after 96-hour induction with doxycycline. **(B)** Expression of DCX and TUBB3 across DIV1-4 in CTRL0 (unedited) and DCX-mTID cell lines, alongside accumulation of biotinylated proteins by Western blot. Knock-in construct is illustrated. **(C)** Immunofluorescence (IF) images show DIV3 neurons with HA staining in CTRL0 (no HA tag) and DCX-mTID (contains HA tag) cell lines (scale bars are 50 µm). Boxed regions are enlarged to emphasize growth cone staining (scale bars are 20 µm). **(D)** IF images show DCX localization in CTRL0 cells fixed at DIV2-5. Scale bars are 100 µm. **(E)** Representative line profiles for fluorescence intensity of MAP2 (magenta), DCX (grey) and TUBB3 (cyan) along neurites at DIV3 and DIV4. Profiles are normalized between channels for length, and within channels according to minimum (background) and maximum (soma) signal. **(F)** Scatter plot showing ratios of MAP2 (magenta) or DCX (grey) intensity to TUBB3 intensity, as a median per neurite (faded circles), per field of view (open circles) and per DIV (black circles).

Differentiation progressed at roughly 24-hour intervals, as visualized by brightfield and/or polarized light microscopy (**Figure 1 supplement 1C, 1D**). DIV2 marked the rough cortical commitment stage, when cells began adopting bipolar orientation. By DIV3 (72 h), cells had extended processes spanning an elongated soma, indicative of nucleokinesis. At DIV4, some cells remained DIV3-like, while the bulk exhibited clustered cell bodies, and processes began to form a mesh-like tissue. Moreover, these soma clusters became denser, and processes became finer and more integrated between DIV4-7, which is generally considered the “maturation” window when axonal-dendritic compartments are constructed and synapses begin to form. Based on their general morphological features, we considered these phases as representative of “progenitor-like” (DIV2), “migration-like” (DIV3), “polarization/onset of maturation” (DIV4), and “maturation” (DIV5) state.

Since DCX’s localization to the growth cone is controlled by phosphorylation of the C-terminal tail (Jin, J. *et al*., 2010), we fixed neurons at DIV3 (migration-like state) to observe DCX localization among growth cone microtubules. Indeed, HA staining indicated the presence of DCX-mTID at the growth cone (**Figure 1C**), allowing us to conclude that incorporation of the tag did not prevent normal localization. Given their synchronized morphological development, we fixed neurons at each DIV (2-5) and stained for MAP2 alongside DCX and TUBB3, to support our classification of morphological staging (**Figure 1D**, **Figure 1 supplement 2**). We found that whereas all three species correlated spatially at DIV3, the normalized signal intensity of MAP2 dropped along neurites at DIV4 and remained low at DIV5 (**Figure 1E, 1F**). Additionally, DCX and TUBB3 localized broadly along neurite projections at DIV3, before becoming enriched periodically along neurites, forming a "dash-like" pattern translated as signal peaks (**Figure 1D-E**, **Figure 1 supplement 2A-D**). This pattern corresponded with DCX’s enrichment of collateral branching sites along the axon, which has been observed in hippocampal neurons (Tint, I. *et al*., 2009). Together, these observations supported our designation of progenitor-like (DIV2), migration-like (DIV3), polarization (DIV4) and maturation (DIV5) as distinct morphological phases of cortical differentiation that constitutes DCX’s native environment.

### DCX undergoes proximity shifts that coincide with differentiation stages

To generate a molecular "atlas" of DCX’s proximal environment across these stages, media was continuously supplemented with biotin to allow for accumulated BioID-labeling throughout differentiation, until cell harvest. Typically, neuronal applications of BioID are limited by their high background (Branon, T. C. *et al*., 2018; Cho, K. F. *et al*., 2020; May, D. G. *et al*., 2020). However, genomic fusion provided us with the opportunity to leverage DCX’s endogenous DIV2 expression onset for quantitative proteomic comparison between DIVs. Biotinylated proteins were identified via liquid chromatography-mass spectrometry (LC-MS) in data-independent acquisition (DIA) mode, which yielded a dataset of 2888 proteins, identified by a minimum of 2 unique peptides and a false-discovery rate (FDR) of 0.5%, for a final full dataset of 2735 proteins (**Figure 2 supplement 1A**). Neurons for BioID harvest were seeded to confluence, to ensure similar cell density for replication and comparison between DIVs. This was supported via principal component analysis (PCA), which confirmed a tight clustering of replicates (**Figure 2 supplement 1B**).

**Figure 2.**
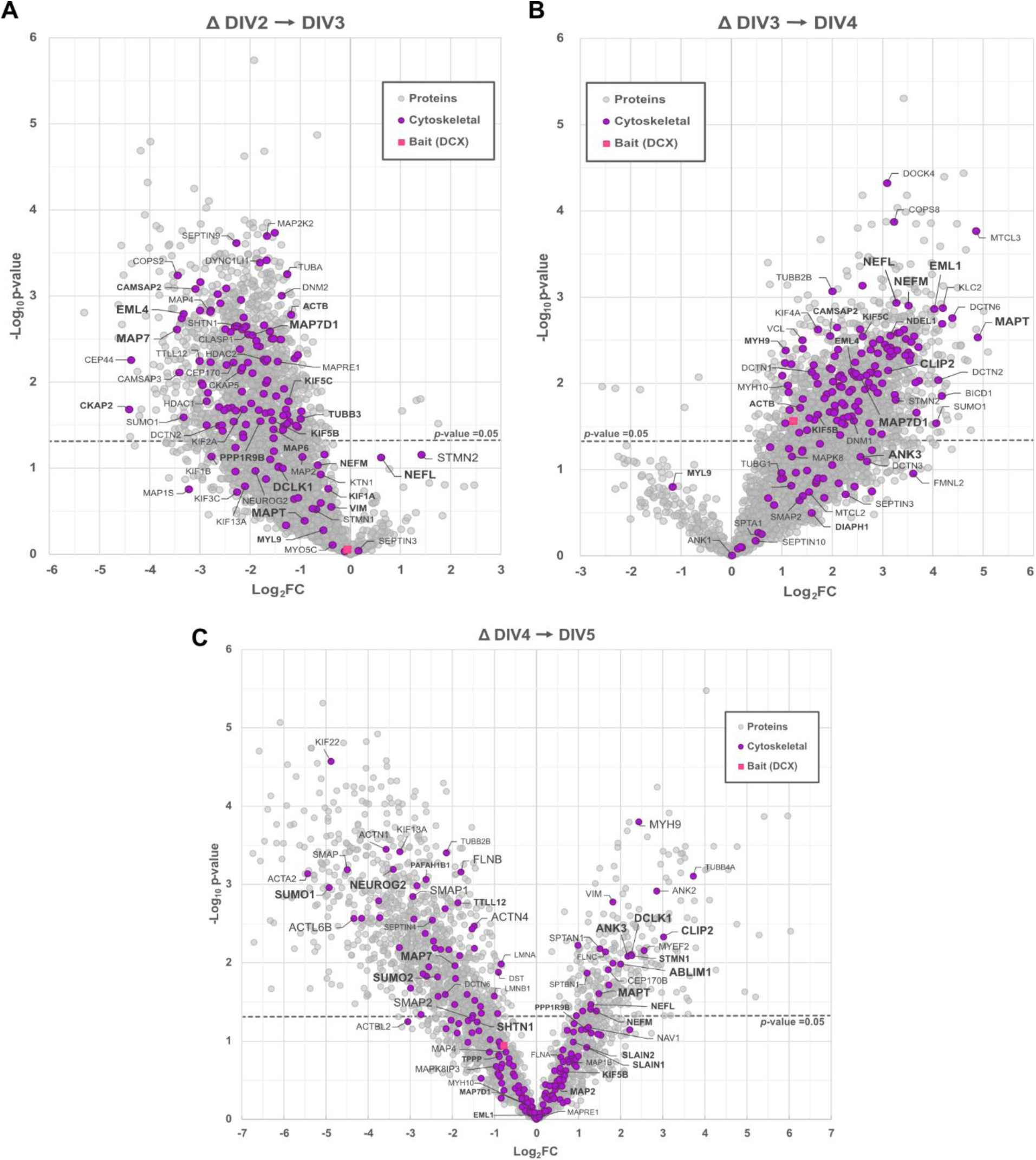
(with 2 supplements) – *The DCX proteome undergoes significant remodeling throughout cortical differentiation*. **(A-C)** Volcano plots illustrate differential enrichment of bulk (grey) and Cytoskeletal (purple) proteins identified by BioID between DIV2–DIV3 (left to right; A), DIV3–DIV4 (left to right; B), and DIV4–DIV5 (left to right; C). Bait (DCX) is indicated in magenta. P-value significance (p<0.05) is indicated by dashed lines. For all plots, n≥3. Plots were generated via Student’s T-test with Benjamini-Hochberg (BH) adjustment on log2-transformed intensity values between DIVs, according to BH-adjusted p-value (-Log10 p-value) and Difference statistics (Log2FC) for each group. Proteins represented by less than 2 valid values in both testing groups were excluded. Remaining invalid values were imputed using a downshift of 1.8 and width of 0.3 standard deviations.

For all species, protein-level intensities were log2-transformed and imputed using standard width (0.3) and downshift (1.8) parameters prior to T-testing (S0=0.1) with the Benjamini-Hochberg (BH) correction for multiple testing (Benjamini, Y. & Hochberg, Y., 1995). BH-adjusted p-values (-log10p-value) and Difference statistical values (Log2FC) were used to construct volcano plots summarizing the enrichment of species between DIV2-3 (**Figure 2A**), DIV3-4 (**Figure 2B**), and DIV4-5 (**Figure 2C**). Surprisingly, volcano plots revealed a global decrease in protein intensity from DIV2-3 (**Figure 2A**), followed by a reciprocal increase from DIV3-4 (**Figure 2B**). This was not due to cell death that typically accompanies DIV3 at lower cell densities, as evidenced by the consistent accumulation of biotinyl species (**Figure 1B**) and the steady detection of the bait (DCX) (**Figure 2A-B**).

To selectively survey DCX’s proximity among cytoskeletal proteins, we used gene ontology (GO) enrichment to parse out known cytoskeletal genes (Combes, F. *et al*., 2021; Locard-Paulet, M. *et al*., 2024). However, these tools identified only a small fraction of proteins (1-2%, ∼35 out of 2735) as cytoskeletal in our bulk dataset. To overcome this limitation, we manually curated a “Cytoskeletal” protein list through cross-referencing of gene IDs with UniProt and Swiss-EMBL designations. While this method was not high-throughput, it allowed us to repeatably parse out ∼10% (250-300 out of 2735) of identified proteins according to their cytoskeletal association. To validate our Cytoskeletal database, we used g:profiler (Kolberg, L. *et al*., 2023) to perform GO over-representation analysis (GO-ORA) and determine the statistical representation of gene terms against a custom background containing the full list of 2735 proteins. As expected, cytoskeletal terms were strongly enriched, with cytoskeletal and microtubule/tubulin-associated terms ranking as most significantly enriched, across Biological Process (BP), Cellular Component (CC), and Molecular Function (MF) categories (**Figure 2 supplement 2**). Highlighting the differential enrichment of cytoskeletal proteins confirmed an increased proximity of maturity-linked proteins from DIV3-4 and DIV4-5, such as tau (*MAPT*), neurofilaments (*NEFM/NEFL*), and ankyrin-G (*ANK3*) (**Figure 2C**) (Barnes, A. P. & Polleux, F., 2009).

Evaluation of Cytoskeletal protein intensities across DIVs revealed that the apparent proximity "dropout" occurring at DIV3 was not simply a turnover event. Rather, it constituted a range of proximal behaviors: some species exhibited temporarily decreased signal at DIV3 which was recovered again at DIV4, whereas others dropped below detection entirely, or remained below detection until DIV4 (**Figure 3A, 3B**). These intensity patterns were not due to protein expression, evidenced by the consistent expression of TRIM46, which dropped out of proximity at DIV3 (**Figure 3C**, **Figure 3 supplement 1A, 1C**), and EB3 (*MAPRE3*), which was not detected at all (**Figure 3 supplement 1A, 1B**). The bulk reduction also contrasted with a cumulative increase of compartment-specific MAPs and neurofilaments (**Figure 3C, 3D**), and the increase of nucleoskeletal components nesprin-2 (*SYNE2*) and SUN1 (**Figure 3D**) (Adam, S. A., 2017; Zhang, X. *et al*., 2009). Heatmaps were also constructed for all other proteins in our Cytoskeletal database to generate a proteomic atlas cataloguing the developmental proximity of cytoskeletal species, using DCX as bait (**Figure 3 supplement 1-5**). While these values are not comparable between species, cumulative labeling and quantitative detection enables DIV-to-DIV comparison, describing the relative proximity of species to DCX across morphological stages. We therefore concluded that, while DCX remains broadly localized throughout development, its local proteomic environment undergoes significant remodeling between morphological stages.

**Figure 3.**
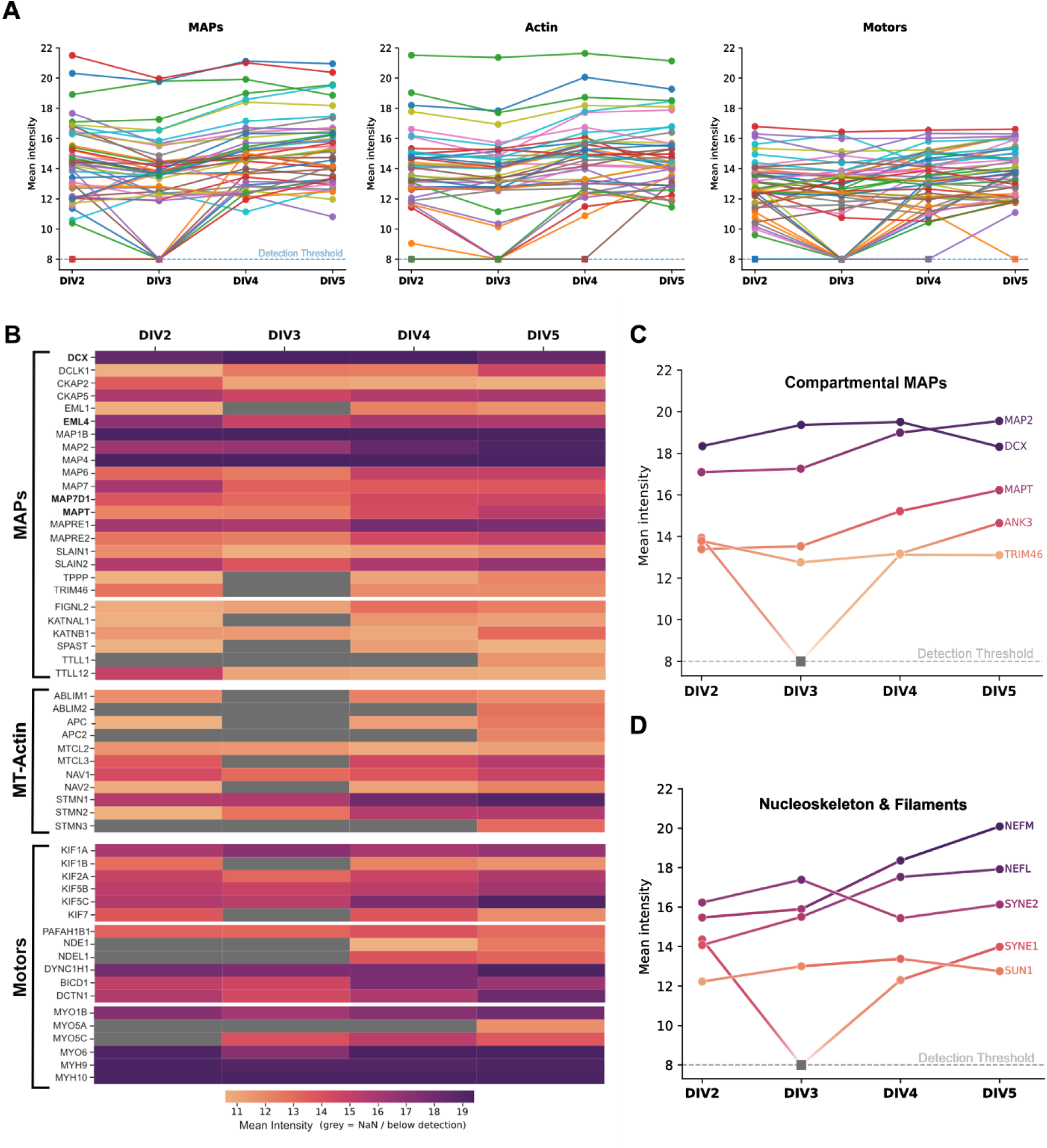
(with 5 supplements) – DCX proximity reveals cytoskeletal remodeling shifts during migration. **(A)** Line plots show protein-level intensity across DIV2-5 for curated Cytoskeletal proteins categorized by UniProt as MAPs, Actin-associated proteins, or motors. Values at or below detection are set to the Detection Threshold marked by a dashed line. **(B)** Representative heat map shows intensity for select Cytoskeletal proteins across DIV2-5. Proteins represented by >2 valid values within a DIV group are shown in grey. **(C-D)** Line plots for select MAPs (C) and Cytoskeletal proteins (D) demonstrate the intensity calibration of heat map color coding, with a dashed line to denote Detection Threshold (∼8). For all panels, n≥3. Summary statistics (mean and standard deviation) were calculated using log10-transformed intensity values within conditions (DIV replicates). Groups with less than 2 valid values were excluded from statistical analysis and marked as “below detection”. Heat map analysis of all proteins in our Cytoskeletal database are catalogued in Figure 3 supplement 1-5.

### R178L mutation reveals a lattice-dependent proximity network

Having characterized DCX’s broad proximal environment, we sought to further interrogate its functional proximity and elucidate networks that are disrupted with DCX mutation. We therefore generated a second iPSC line to express the R178L mutation fused with miniTurboID (R178L-mTID). R178L is directly implicated in lissencephaly (Des Portes, V. *et al*., 1998b), without a clear explanation for its effect on migration. Indeed, whereas other mutants exhibit structural instability, disrupted binding, and/or localization defects (Bechstedt, S. *et al*., 2012; Gleeson, J. G. *et al*., 1999b; Yap, C. C. *et al*., 2016), R178L exhibits wildtype-like expression, localization and phenotypic markers (**Figure 4B**, **Figure 4 supplement 1A-B**), with the exception that it does not exhibit the same sigmoidal binding behavior suggestive of cooperativity *in vitro* (Bechstedt, S. *et al*., 2012). We postulated that R178L would expose disease-relevant MAP networks which are sensitive to DCX’s cooperatively (stably) bound state (**Figure 4A**). BioID samples for both DCX(WT)-mTID and DCX(R178L)-mTID were harvested at DIV5 to emphasize cumulative labeling of molecular neighbours. DIA LC-MS revealed that within sample batches, replicates clustered very tightly (*n*=4 for both conditions; **Figure 4 supplement 2A, 2B**), while increased variance between batches (*N*=2) minimized the appearance of imputation-based artifacts, which was prevalent among species with intensities near the detection floor.

**Figure 4.**
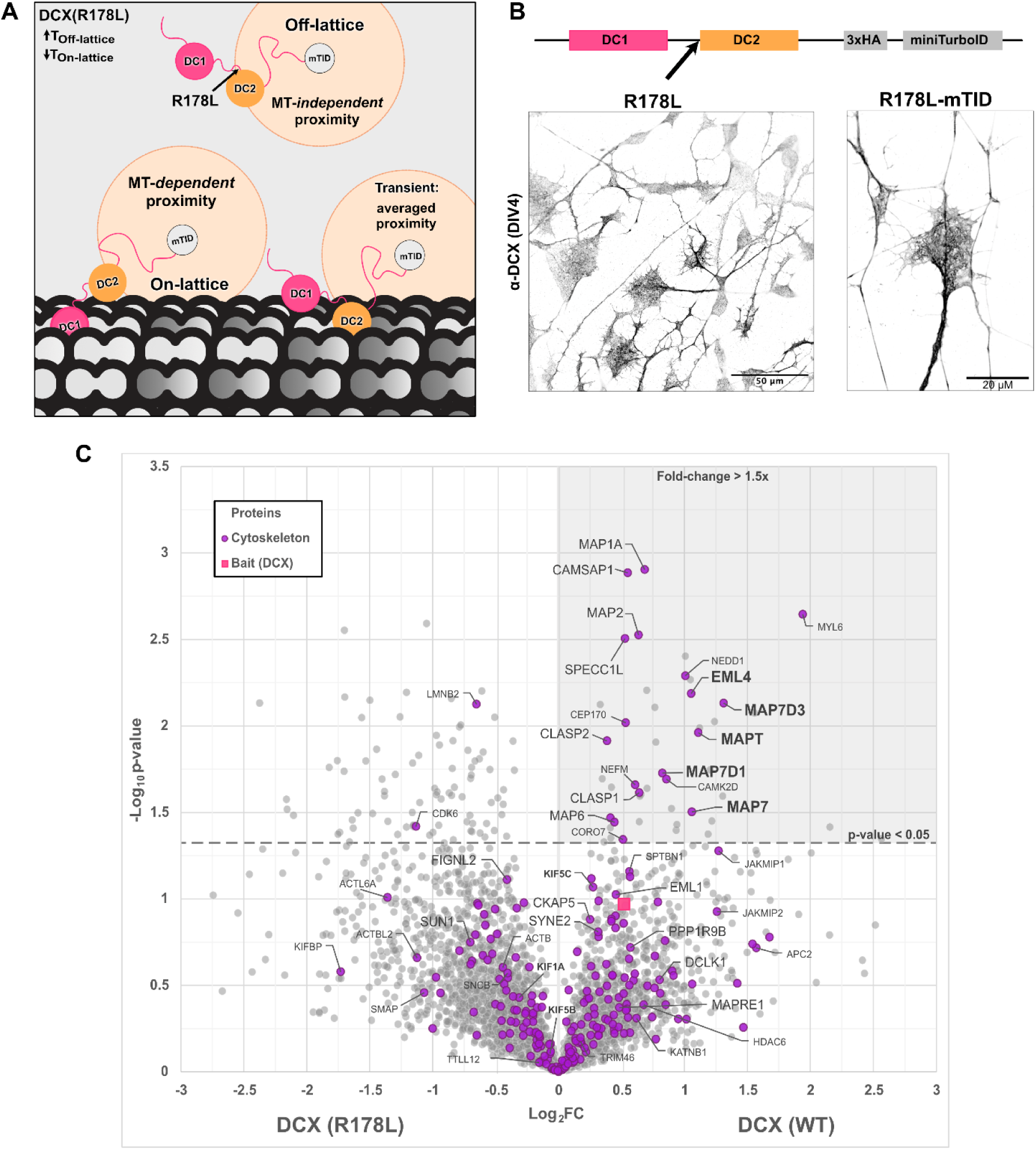
(with 2 supplements) – R178L mutation reveals a lattice-dependent proximity network. **(A)** Conceptual schematic illustrates predicted redistribution of biotin-labeling caused by p.Arg178Leu (R178L) mutation. **(B)** Immunofluorescence images show DIV4 localization of untagged R178L (left; scale bar is 50 µm) and growth cone localization of R178L-mTID (right; scale bar is 20 µm). **(C)** Volcano plot shows differential enrichment of bulk (grey) and cytoskeletal (purple) proteins identified by BioID in DCX-mTID (WT) and R178L-mTID (R178L) neurons at DIV5. Bait (DCX) is indicated in magenta. The shaded area indicates statistical and enrichment significance (p-value <0.05; fold-enrichment >1.0x). Statistical significance was determined via Student’s t-testing (S0 = 0.1) with BH-adjustment). For each condition, n=4. Proteins with less than 3 valid values in at least one group were excluded from statistical testing, and remaining missing values were imputed using standard downshift (1.8) and width (0.3) parameters.

To identify proteins significantly enriched in the WT condition, we constructed a volcano plot using BH-adjusted -log10p-values and Difference statistics with positive Log2FC values representing WT-enrichment (**Figure 4C**). Consistent with our original hypothesis, GO-ORA analysis confirmed that WT-enriched species were significantly over-represented for “tubulin binding” and “microtubule binding” MF terms, against the full 2735-protein background (**Figure 5 supplement 1**). These results were a low estimate, since enriched candidates such as EML4, MAP7, and MAP7D1 are not yet annotated for either of these terms, while all three are known MAPs with experimentally determined microtubule-binding activity (Hooikaas, P. J. *et al*., 2019; Houtman, S. H. *et al*., 2007; Pollmann, M. *et al*., 2006).

**Figure 5.**
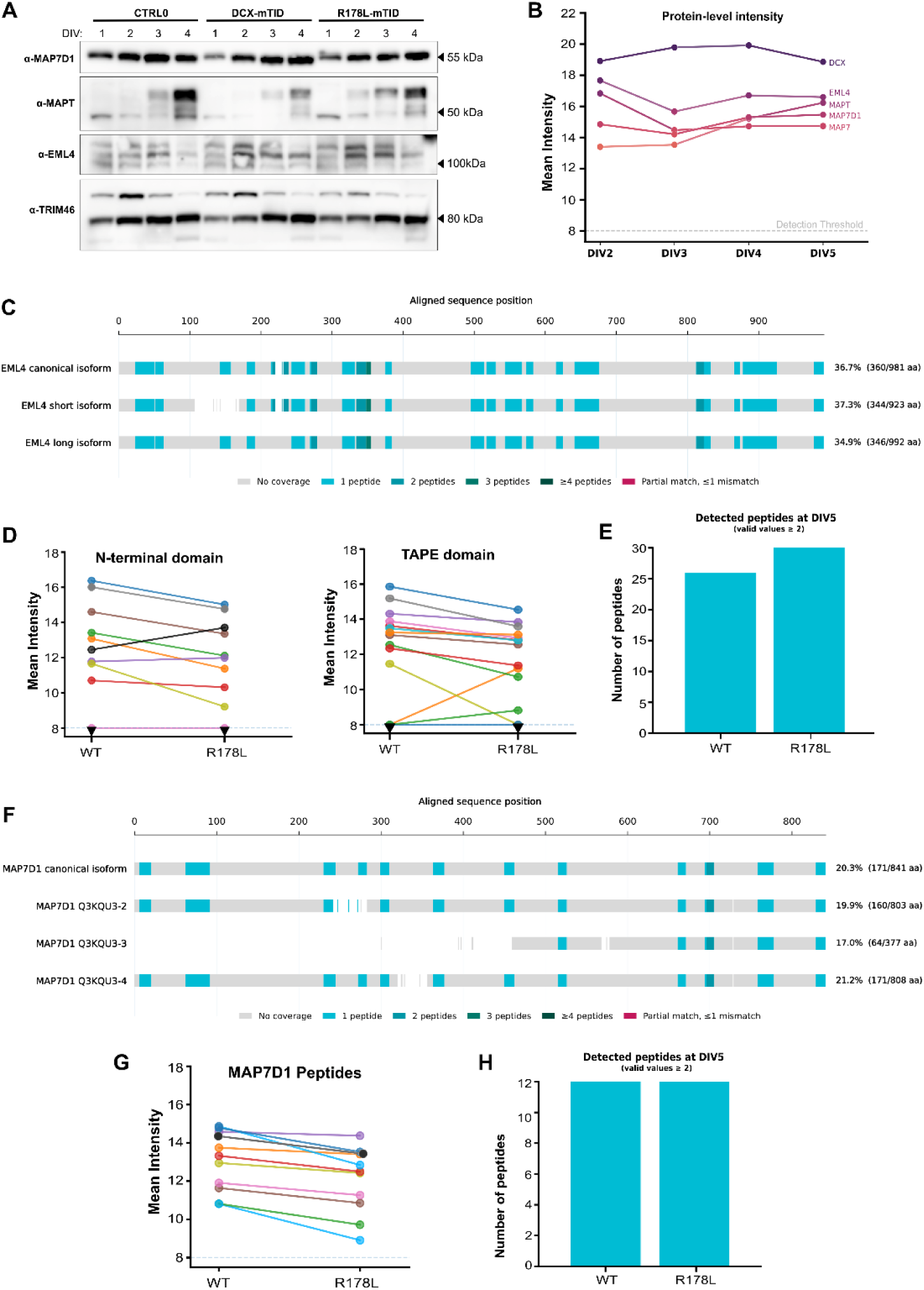
(with 3 supplements) – *WT-enrichment of EML4 and MAP7D1 is reflected in protein- and peptide-level intensities*. **(A)** Immunoblot validation of selected enriched proteins (MAP7D1, MAPT, and EML4) across DIV1-4 for CTRL0, DCX-mTID and R178L-mTID cell lines. TRIM46 is shown as a developmental control. For each species, n≥3. **(B)** Line plot shows intensity of selected enriched proteins, cross-referenced with data obtained in Figure 3. **(C, F)** Multiple sequence-alignment of Uniprot-annotated isoforms of EML4 (C) and MAP7D1 (F). Peptides detected via LC-MS are indicated in teal, with darker shades indicating overlap of distinct peptides. All peptides were mapped to at least one isoform. **(D, G)** Pairwise comparison of peptide intensities detected for EML4 (D) and MAP7D1 (G) between DCX(WT) and DCX(R178L). **(E, H)** Bar chart illustrates the number of peptides detected between WT and R178L conditions for EML4 (E) and MAP7D1(H). Peptide intensities represented by less than 2 valid values were considered below detection.

Given the change-of-function of R178L (as opposed to loss-of-function), enrichments were slight. To ensure enrichments were not biased by differential expression between cell lines, we performed immunoblotting for select species: MAP7D1, EML4, and MAPT (tau). Between DCX-mTID and R178L-mTID, all three MAPs exhibited similar expression patterns (**Figure 5A**) and a consistent proximity to DCX across developmental stages (**Figure 5B**). This was apparently true of all significantly enriched (p-value < 0.05, FC >1.0) species according to protein-level intensities (**Figure 5 supplement 3A**). However, some of these proteins were represented by only a few peptides in either condition (**Figure 5 supplement 2**). The R178L-sensitivity of MAP7D1 and EML4 was intriguing, since both have been captured in co-immunoprecipitation studies of DCLK1 (DCX superfamily kinase) (Koizumi, H. *et al*., 2017), but not DCX (Fu, X. *et al*., 2022). EML4 exhibited a robust DCX proximity throughout differentiation, both at the protein- and peptide-level (**Figure 5B**, **Figure 5 supplement 3B**), with peptides mapping to >30% sequence coverage (**Figure 5C**). Moreover, its ∼2-fold WT-enrichment was not due to a decreased in the number of detected peptides in the R178L condition (**Figure 5E**). Rather, most R178L-detected peptides exhibited reduced intensities than in WT (**Figure 5D**), suggesting that in DCX-mTID samples, DCX spends slightly more time around EML4 than R178L-mTID. MAP7D1 peptides exhibited a similar pattern, though not with the same robustness as EML4 (**Figure 5F-H**).

### MAP7D1 and EML4 partially correlate with DCX but enrich distinct compartmental networks

Since EML4 and MAP7D1 bind and stabilize microtubules (Kikuchi, K. *et al*., 2022; Pollmann, M. *et al*., 2006), we sought to visualize the spatial proximity of both MAPs in relation to DCX/TUBB3. Consistent with previous reports, MAP7D1 enriched the soma and neurites, while avoiding the growth cone and collateral branching sites across DIVs, while DCX remained broadly localized (**Figure 6A**) (Koizumi, H. *et al*., 2017). Moreover, MAP7D1 signal along neurites underwent a shift in process selectivity, enriching all processes at DIV3 but only enriching a subset at DIV4-5 (**Figure 6A, 6C, Figure 6 supplement 1**). Conversely, EML4 localized more broadly with DCX than MAP7D1, though with greater enrichment covering the entirety of the soma. EML4 also remained correlated (*r* >0.6) with TUBB3/DCX from DIV3-4, based on spatial overlap (**Figure 6 supplement 2E**) and normalized signal intensity (**Figure 6C-D**) measured along neuronal processes. While measurements emphasized the overlap of all four species along neurites, enrichment patterns within the soma revealed clear divergence between them. Specifically, DCX/TUBB3 consistently enriched regions in front and behind the elongated nucleus at DIV3, before becoming polarized as soma clustered, between DIV4-5 (**Figure 6 supplement 1B, 2B**). In comparison, EML4 enriched the soma almost entirely, surrounding the nucleus, while MAP7D1 enriched a centrosomal pattern similar to Lis1 (Tanaka, T. *et al*., 2004), consistent with previous reports (Koizumi, H. *et al*., 2017). Therefore, while these results support our findings that EML4 and MAP7D1 exhibit significant compartmental proximity with DCX, their enrichment patterns are distinctly altered as neurons migrate and mature.

**Figure 6.**
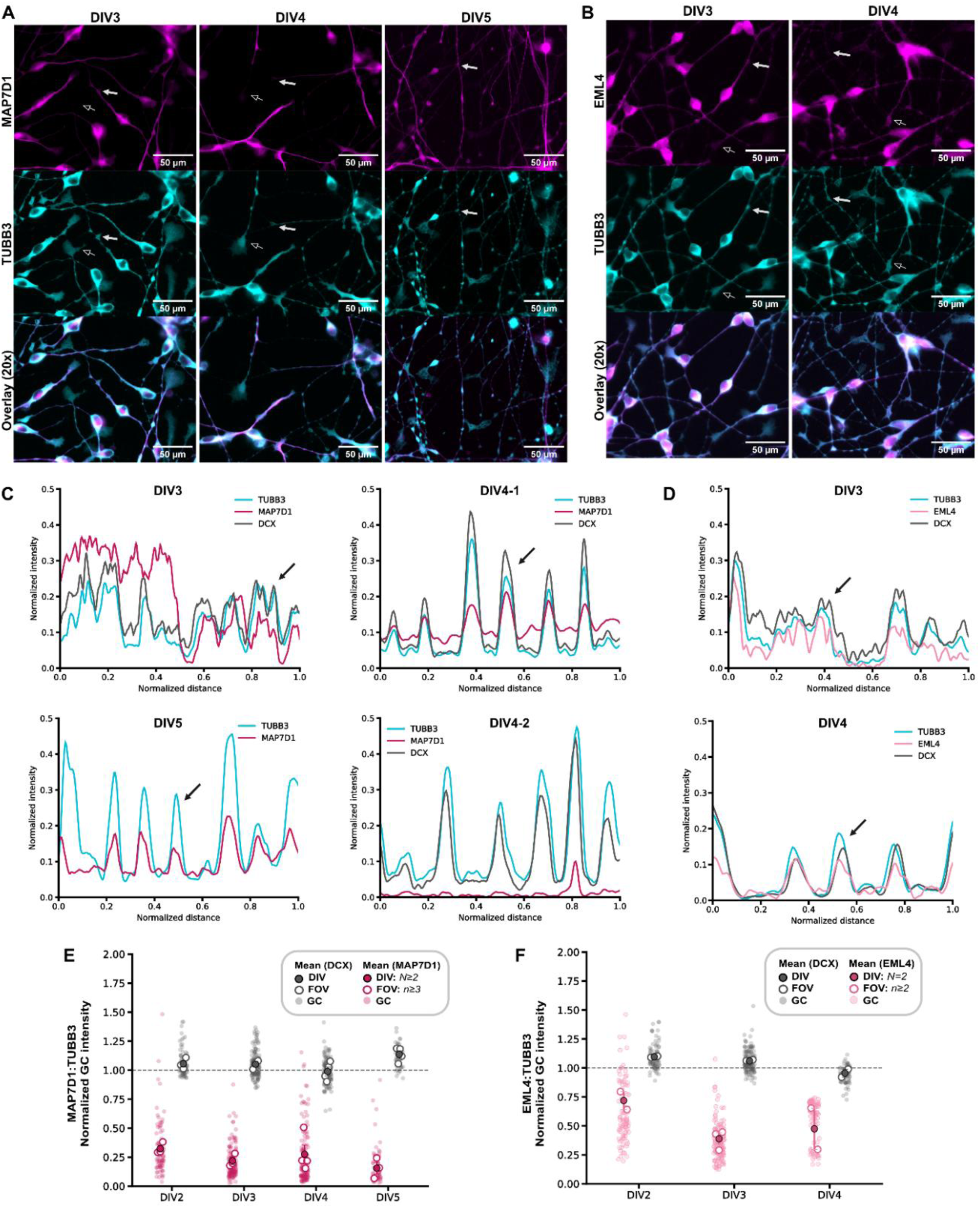
(with 2 supplements) – *MAP7D1 and EML4 partially correlate with DCX but enrich distinct microtubule networks*. **(A-B)** Widefield images of neurons stained for MAP7D1 (magenta) at DIV2-5 (A) or EML4 (pink) at DIV3-4 (B), along with TUBB3 (cyan). Growth cones are denoted by open arrows; branch junction sites are denoted by solid arrows. Scale bars are 50 µm. **(C-D)** Representative fluorescence profiles for DCX (grey), TUBB3 (cyan), and MAP7D1 (magenta; C panels) or EML4 (pink; D panels) along neurites at each DIV. **(E-F)** Scatter plots show normalized growth cone intensity ratio of DCX (grey) and either MAP7D1 (magenta) or EML4 (pink) to TUBB3 at each DIV. Data are presented as the mean intensity per ROI (GC; faded circles), per field of view (open circles) and per DIV (black circles); data represented by a minimum of 2 biological replicates, 2 distinct fields per sample.

## Discussion

In this work, we sought to elucidate the endogenous molecular context of DCX, which has been challenging to characterize. Until recently, studies involving immature neuronal tissues have been limited to primary neurons and heterogeneous culture tissues (Fu, X. *et al*., 2022; Wong, H. H.-W. *et al*., 2023). Our data highlight this limitation, finding that distinct phases are characterized by bulk proteome reorganization (**Figure 2**, **Figure 3**). Proteins blotted across DIV1-5 exhibited band patterns consistent with alternative induction models that take place over a 30-day transition from stem cell to neural progenitor, followed by a 7-day terminal differentiation (**Figure 1B, 6A**) (Madgwick, A. *et al*., 2015; Sébastien, M. *et al*., 2025). EML4, specifically, undergoes a distinct splicing shift visible in our NGN2-directed, 96-hour differentiation as in the terminal differentiation period of a previous model (**Figure 5A**) (Madgwick, A. *et al*., 2015). In addition, MAP7D1 adopts a localization in our differentiating cells that is consistent with cultured hippocampal neurons, where it likewise avoids the DCX/DCLK1-enriched growth cone (**Figure 6A**, **Figure 6 supplement 1A, 1B**) (Koizumi, H. *et al*., 2017). These independent observations provide confidence that our proteomic findings are not specific to NGN2-specific pathways but may represent broad cytoskeletal rearrangements that widely occur during the terminal differentiation process. NGN2-directed induction therefore enabled visualization of these trends and will be a valuable tool for elucidating the molecular underpinnings of developmental processes in the future.

One possible explanation for the DIV3 proximity shift was DCX’s reported enrichment at the growth cone (Dema, A. *et al*., 2024), given the similar knock-in strategy and tag placement. However, this hypothesis was inconsistent with our earlier observations that DCX localized broadly with TUBB3 throughout differentiation (**Figure 1C, 1D, Figure 1 supplement 2B**). We considered this discrepancy might arise due to kinetic bias between live-cell and fixed-cell imaging, since *in vitro* reconstitutions suggest DCX can bind microtubules with variable affinity depending on its mode of engagement (Bechstedt, S. *et al*., 2012; Bechstedt, S. *et al*., 2014; Manka, S. W. *et al*., 2020). Thus, whereas cell fixation may account for microtubule-bound DCX regardless of its affinity or dynamic engagement, live-cell imaging may emphasize regions of dynamic engagement or high enrichment (Irgen-Gioro, S. *et al*., 2022). The reduced expression and specific targeting of DCX isoform d (transcript 5; NM_001195553.2), which was consistent between the two studies (Dema, A. *et al*., 2024), was not responsible for this bias, since both edited (DCX-mTID) and unedited (CTRL0) cell lines exhibited broad distribution of DCX, by antibodies against both DCX and the HA tag (**Figure 1C, 1D**).

Instead, we considered that whereas DCX remains tightly correlated with TUBB3 in various compartments, there are significant morphological differences between DIV2-3, as well as DIV3-4 which could be responsible for observed proximity shifts. For example, DCX is implicated in the coupling between centrosomal and perinuclear (acentrosomal) microtubules during nucleokinesis (Tanaka, T. *et al*., 2004; Vinopal, S. & Bradke, F., 2025), which occurs between DIV2-4 (**Figure 1D**, **Figure 7A, 7B**). In contrast, DIV4 marks the relocalization of DCX (and TUBB3) to periodic sites along processes which become extensively branched by DIV5 (**Figure 1D**, **Figure 1 supplement 2, Figure 6A**, **Figure 6 supplement 1**). This pattern is consistent with implications for DCX in collateral branching of the axon (Kappeler, C. *et al*., 2006; Tint, I. *et al*., 2009), a process which occurs after polarization and requires a host of other cytoskeletal proteins. Interestingly, several of these proteins are WT-enriched relative to the R178L mutation, including CEP170, CORO7, NEFM/neurofilaments, MYL6+SPECC1L, and NEDD1 (**Figure 4C**, **Figure 5 supplement 2**) (Berger, S. L. *et al*., 2018; Burnette, D. T. *et al*., 2008; Kevenaar, J. T. & Hoogenraad, C. C., 2015; Liao, Y.-C. *et al*., 2026; Saadi, I. *et al*., 2023; Shatery Nejad, N. *et al*., 2025; Vinopal, S. *et al*., 2023). Moreover, many of these maintain a high level of proximity to DCX throughout differentiation (**Figure 3B**, **Figure 3 supplement 1-5, Figure 5 supplement 3A**). Given the implications for DCX in both nucleokinesis and axonal branching (Tanaka, T. *et al*., 2004; Tint, I. *et al*., 2009), it is possible that DCX operates in a similar role – linking centrosomal and acentrosomal microtubule populations – to support both developmental activities as distinct spatiotemporal “programs” (**Figure 7**). Together with DCX’s reported activities at both migration and axonal growth cones (Bott, C. J. *et al*., 2020; Dema, A. *et al*., 2024) and within dendritic compartments (Li, P. *et al*., 2021), the proximity decrease observed between DIV2-4 likely does not represent DCX sequestration from many species. Rather, our findings likely reflect the global reorganization of the neuronal cytoskeleton as microtubule populations become more functionally diverse.

**Figure 7.**
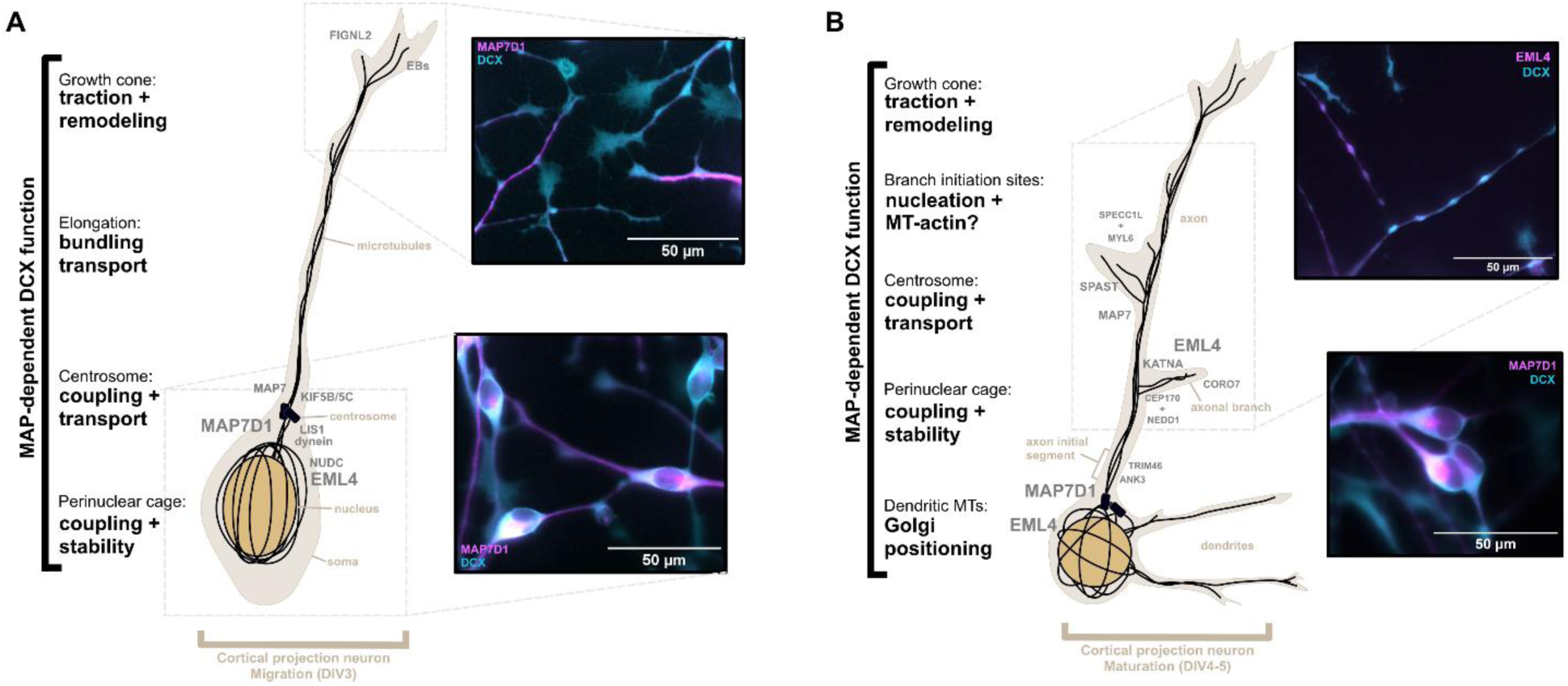
Hypothetical model for DCX bridging distinct compartmental networks. **(A-B)** Integrative schematic illustrating the proposed functions for DCX depending on compartmental MAP enrichment that varies with morphological staging. Illustrations depict experimentally determined and hypothetical enrichment patterns in migrating and maturing neuronal cultures for MAPs which exhibit proximity to DCX (Figure 2-4, Figure 3 supplement 1-5). Inset micrographs show overlay of DCX (cyan) and either MAP7D1 or EML4 (magenta). Micrograph scale bars are 50 µm.

While we specifically set out to elucidate which cytoskeletal proteins share lattice space with DCX and populate its endogenous environment, we were intrigued by R178L’s ability to highlight MAP networks with apparent functional coherence (**Figure 4C**, **Figure 7**). Of note, the proteins screened and identified here do not necessarily interact with DCX. However, EML4, MAP7D1, MAPT, MAP2, DCX, and SPECC1L, among others identified, all stabilize microtubules through distinct mechanisms (Dehmelt, L. & Halpain, S., 2004; Kikuchi, K. *et al*., 2022; Moores, C. A. *et al*., 2004; Pollmann, M. *et al*., 2006; Saadi, I. *et al*., 2023). Moreover, their distinct mechanisms intersect with other functional activities: MAP7D1 stabilizes acetylated microtubules, while supporting MAP7-directed kinesin-1 activation (Hooikaas, P. J. *et al*., 2019; Kikuchi, K. *et al*., 2022), and MAP2/MAPT stabilize microtubules and prevent microtubule-severing by katanin (*KATNA, KATNB, KATNAL)* (Qiang, L. *et al*., 2006). Meanwhile, MAP2 empties from axonal compartments (**Figure 1E-F**, **Figure 1 supplement 2A, 2C**) (Caceres, A. *et al*., 1984) at the same time that tubulin becomes heavily polyglutamylated and spastin (*SPAST*) severing enzymes become detectable in DCX proximity (**Figure 3B**, **Figure 4 supplement 1A, 1B**) (Gavoci, A. *et al*., 2025; Kuo, Y.-W. & Howard, J., 2021; Yu, W. *et al*., 2008). Thus, rather than identifying which proteins interact meaningfully with DCX, our proximity perturbation by R178L revealed a functional convergence among MAPs that may confer both synergistic and specific regulatory activities, depending on developmental timing.

Together, our findings present both a spatiotemporal and functional cytoskeletal context for DCX. Considering observations that MAPs compete for lattice space and influence over polymer dynamics and intracellular transport (Dema, A. *et al*., 2023; Dema, A. *et al*., 2024; Monroy, B. Y. *et al*., 2018; Monroy, B. Y. *et al*., 2020), studies identifying protein neighbours with functional symmetry (that are not necessarily interaction-based) are increasingly important. For example, the identification of three MAP7-domain-containing family members in a DCX proximity screen may seem paradoxical, given their preference for distinct microtubule conformations and oppositional influence over kinesin-based transport. However, our results don’t establish that these proteins interact. Rather, R178L perturbation reduces proximity to these proteins, providing a cellular context in which their lattice preferences and regulatory activities can be interpreted. For example, we show that DCX and TUBB3 are tightly spatially coupled (**Figure 1E-F**, **Figure 1 supplement 2, Figure 6C-D**, **Figure 6 supplement 1-2**), echoing previous studies that emphasize their phenotypic and developmental implications (Poirier, K. *et al*., 2010; Qu, C. *et al*., 2013; Shao, Q. *et al*., 2019). Meanwhile, this is not true of DCX and pan-α-tubulin antibodies like DM1α (Tint, I. *et al*., 2009). We also find that MAP7D1 and EML4 overlap with (but are not restricted to) DCX/TUBB3-enriched regions (**Figure 6A-B**, **Figure 6 supplement 1-2**). Thus, the specific microtubule-localization of these proteins may reflect features of the tubulin code (Janke, C. & Kneussel, M., 2010; Verhey, K. J. & Gaertig, J., 2007), with distinct microtubules being bundled or coupled together. Our findings therefore provide proteomic constraints and variables for the development of future reconstitution approaches. Together with cellular assays, proximity-based approaches provide a proteomic lens to aid in the interpretation of biophysical phenomena and advance our understanding of cytoskeletal coordination.

## Materials and Methods

### Generation of CRISPR-Edited iPSC Lines

Human induced pluripotent stem cells (iPSCs) containing a doxycycline-inducible NGN2 promoter (European Bank for Induced Pluripotent Stem Cells, #BiONi-010-C13) were used as the parental, unedited (CTRL0) line. Cells were subsequently edited via CRISPR-Cas9 to engineer the DCX variant NM_001195553.2:c.[533_534delinsTA;558C>G], encoding p.Arg178Leu (R178L) and a silent p.Arg186= substitution introduced to disrupt the Cas9 protospacer adjacent motif (PAM) sequence. Sequencing information is provided in **Supplemental File 26**.

To generate the R178L cell line, iPSCs were resuspended in a DPBS-based buffer and transfected with pX458 plasmid encoding spCas9 and gRNA sequences targeting the R178 locus and single-stranded DNA (ssDNA) oligonucleotides containing nucleotide substitutions. Cells were transfected using a single electrical pulse of 65 V over a 5 ms pulse width, as described previously (Sébastien, M. *et al*., 2025).

To engineer C-terminal miniTurboID-fused cell lines, a double-stranded DNA (dsDNA) donor cassette was synthesized as an E-block gene fragment (Integrated DNA Technologies; sequence provided in **Supplemental File 23**). This homology-directed repair (HDR) template comprised a 5′ homology arm spanning the final 30 base pairs (bp) of the DCX transcript 5 mRNA (NM_001195553.2), an in-frame sequence encoding an HA-tagged miniTurboID ligase flanked by flexible peptide linkers, and a 3′ homology arm spanning 30 bp inclusive of the endogenous transcript 5 STOP codon. Synthetically modified guide RNA (sequence: CGCUUGGUGAUUCCAUGUAA) (purchased from Synthego) was complexed with purified spCas9 nuclease at a ratio of 7.5(300 pmol):1 (40 pmol) to form ribonucleoprotein (RNP) complexes before addition of 2 μL of the dsDNA template (200-300 ng/µL). Aside from the addition of dsDNA template, cells were transfected according to the Synthego IPS Cell Nucleofection Protocol (available at www.synthego.com/resources) with RNP-donor mixture using a Lonza 4D-Nucleofector X Unit and the P3 Primary Cell Kit using program CA137. Nucleofected cells were allowed 5 days to recover and split with Accutase (Stemcell Technologies) for single-cell sorting via a BD FACS Aria III flow cytometer (BD Biosciences). Surviving monoclones (40% survival rate for both WT and mutant) were allowed 14 days recovery prior to sequencing and expansion. To screen for tag incorporation, clones were clump-split with Gentle Cell Dissociation Reagent (GCDR; Stemcell Technologies). A portion of the cell suspension was mixed with QuickExtract DNA extraction solution (Millipore Sigma) and prepared according to manufacturer’s instructions. Genomic DNA was PCR amplified using primers flanking the insert site prior to agarose gel screening. Clones positive for the insert were submitted for sequence analysis via long-read Oxford nanopore sequencing (Plasmidsaurus). Sequence-verified clones were subsequently expanded for storage. Primers and DNA sequences are provided aligned with DCX FASTA in **Supplemental Files 24-26**.

### Cell Culture Maintenance and Neuronal Differentiation

Parental and edited iPSC lines were maintained under feeder-free conditions in mTeSR Plus medium (Stemcell Technologies) on Matrigel-coated substrates (Corning) and routinely passaged using GCDR. Cells were monitored to ensure a low passage number (<10) and routinely screened for mycoplasma contamination. To induce cortical differentiation, iPSCs were dissociated with Accutase and seeded onto dishes pre-coated with poly-L-ornithine (100 ng/mL) and laminin (1 μg/mL; Millipore Sigma). Cortical differentiation was induced by seeding cells into Induction media: BrainPhys Neuronal induction Medium supplemented with SM1 and N2A, dibutyryl-cAMP, ascorbic acid, brain-derived neurotrophic factor (BDNF), glial cell line-derived neurotrophic factor (GDNF), doxycycline (2 μg/mL), and 50 μM d-biotin (concentrations and supplier information is provided in Key Resources Table). Induction media was replenished daily, with doxycycline being omitted after DIV4, according to extended induction protocols (Muñoz-Estrada, J. *et al*., 2026). The addition of Rho-kinase (ROCK) inhibitor Y-27632 was restricted to post-thaw recovery, post-electroporation survival, and low-density single-cell seeding steps required for imaging analysis.

### Lysate preparation and Western blotting

Cells were seeded to confluence in 6 cm dishes and harvested within an hour of the 24-hour morphological time points. Prior to harvest, cells were washed twice with warm DPBS, followed by a wash with ice-cold DPBS to encourage cell detachment. Harvested cells were pelleted and resuspended in lysis buffer (50 mM Tris-HCl, 150 mM NaCl, 1 mM EDTA and 1% Triton X-100, pH 7.4) supplemented with Complete Protease Inhibitor (Roche) and PhosSTOP phosphatase inhibitor (Millipore Sigma). Lysates were sonicated prior to flash-freezing of aliquots and were used only once after thaw. Blotting membranes (PVDF) were incubated with 0.1% Ponceau-S staining solution immediately following transfer to confirm equal protein-loading across lanes, with subsequent blocking in TBS supplemented with 0.2% Tween-20 and 5% BSA (3% BSA for antibody incubations). Primary antibodies were incubated with membranes overnight at 4 °C with gentle agitation, followed by 90 minutes incubation with HRP-linked secondary antibodies. Membranes were imaged using an AI600 gel imager (GE Healthcare), to acquire hybrid images with and without pre-stained molecular weight markers. Membranes incubated with proximity-identified species (EML4, MAP7D1, TRIM46) were stripped gently (0.2 M glycine supplemented with 0.1% SDS, 1% Tween-20, pH 2.2) for 10 minutes and re-blocked for incubation with either DCX or TUBB3 (as appropriate) for quality control. Uncropped original and replicate blot images, including corresponding Ponceau-S-stained membrane and molecular weight images are provided as **Supplemental file 28**.

### Proximity Labeling and Streptavidin Affinity Purification

MiniTurboID-expressing cell lines (DCX-mTID/R178L-mTID) were seeded to confluence in 10 cm dishes and harvested at 24-hour intervals corresponding to DIV2, 3, 4, and 5. For each biological replicate, two confluent 10 cm dishes were gently rinsed with DPBS (37 °C) and lysed in 1 mL of ice-cold RIPA buffer (50mM Tris-HCl, 150mM NaCl, 1% NP-40/IGEPAL, 0.5% sodium deoxycholate, 0.1% SDS, 1mM EDTA, pH 7.4) supplemented with protease and phosphatase inhibitor. Lysates were sonicated and clarified via centrifugation at 13, 000 RCF for 10 minutes at 4 °C, and supernatants were pre-cleared by incubation with Sepharose 4B beads to eliminate non-specific resin binders. Pre-cleared supernatants were incubated with Streptavidin-linked agarose beads (Thermo Fisher Scientific) with gentle agitation at 4 °C for 4 hours. Beads were washed extensively with RIPA (no deoxycholate) and flash-frozen in 50 µL of 50 mM ammonium bicarbonate supplemented with 50 µM free d-biotin.

### Mass Spectrometry and Quantitative Proteomics

Biotinylated proteins were eluted from beads with Laemmli buffer and resolved on a stacking gel, followed by in-gel reduction, alkylation with iodoacetic acid, and tryptic digestion. Extracted peptides were solubilized in 0.1% aqueous formic acid.

LC–MS/MS analysis was performed on a Thermo FAIMS Duo Pro/Orbitrap Astral mass spectrometer system coupled to a Thermo Scientific Vanquish Neo UHPLC using mobile phase A: 0.1% formic acid in water and mobile phase B: 0.1% formic acid 80% acetonitrile in water. Briefly, 200 ng of peptide/sample were loaded onto a Thermo PEPMAP Neo 300 μm × 5 mm trap column into an IonOpticks Aurora Elite 150 μm × 8 cm separation column heated to 50C using a 180SPD gradient (4-40% mobile phase B over 7 minutes + wash + reconditioning steps). On the Orbitrap Astral mass spectrometer, a FAIMS voltage of -45V (to emphasize 2+ m/z) was used. MS1 spectra were recorded using the Orbitrap analyzer at a resolution of 240,000 from m/z 380 to 980 using an automated gate control (AGC) target of 500% and a maximum injection time of 3 ms. For MS2 in nDIA mode using the Astral analyzer, 200 nonoverlapping isolation windows from m/z 380 to 980 of 3 m/z were used. HCD collision energy is set to 25% and the RF lens at 40%.

Raw DIA files were searched against the 2025 Human FASTA proteome supplemented with common contaminants and decoys via FragPipe v24.0 using the DIA_SpecLib_Quant workflow. MSFragger-DIA was used for direct identification from DIA spectra, followed by peptide-spectrum match rescoring and FDR filtering. An experiment-specific spectral library was generated from DIA files and used for quantification with DIA-NN v1.8.2 beta 8. Searches permitted two missed tryptic cleavages and included carboxymethylation of cysteine (+58.0055 Da) as a fixed modification. Peptide and protein identifications were initially controlled at a 1% FDR during FragPipe/DIA-NN processing. Raw files, spectral reports and output files are available via the PRIDE ProteomeXchange repository with accession number PDX083262.

### Proteomic Data Processing

Peptide- and protein-level intensity reports generated by FragPipe were imported into Perseus (Max Planck Institute of Biochemistry) for statistical processing, with proteins requiring detection by a minimum of two unique peptides. Sample quality was determined via replication analysis, using hierarchical clustering and Principal Component Analysis (PCA) performed on log2-transformed values (**Figure 2 supplement 1, Figure 4 supplement 2**). PCA required imputation of invalid values using a width and downshift of 0.3 and 1.8 standard deviations, respectively. Replicates displaying distinct batch-dependent clustering were retained to incorporate inter-batch variance.

Volcano plots were generated using pairwise differential enrichment between conditions via Student’s T-testing with Benjamini-Hochberg (BH) adjustment (S^0^ = 0.1) and an FDR threshold of 0.5%. A minimum of three biological replicates was required for repeatability, and proteins with less than 2 valid values in at least one group were excluded from pairwise analysis. To account for differences in expression and proximity between DIVs, filtering for valid values was performed on individual matrices split from the original (comprehensive) matrix. Remaining invalid values were imputed using the standard width (0.3) and downshift (1.8) parameters. Heatmaps and line plots were constructed using summary statistics, to include all proteins. Species represented by fewer than 2 valid values in a particular group/condition were considered “below detection”. For plot generation, Perseus Master files (Supplemental files 2-5) were used to perform statistical analysis, and matrices containing statistical values were exported and converted into either Excel workbooks (.xlsx) files for Volcano plot construction (Supplemental files 6-9), or .csv files (Supplemental files 10-13 and 18-20) for plotting with custom Python scripts. Jupyter Notebooks containing custom analysis and plotting scripts are publicly available in the DCX-proximity GitHub repository.

### Database curation and validation (Gene ontology analysis)

To generate a database of Cytoskeletal proteins from list of 2735 identified proteins, gene names were cross-referenced against UniProt and Swiss-EMBL databases for cytoskeletal association. Specifically, proteins were included based on their association with (or regulation of) microtubules, which included adjacent protein networks such as actin, intermediate filaments, septins, spectrins, regulatory proteins, motor or cargo components. Proteins were parsed manually from the protein-level master Perseus file and split from the main matrix to generate a master list of “Cytoskeletal” proteins (**Supplemental file 2**). To validate the database, we submitted the list of parsed proteins for GO analysis using g:profiler (Kolberg, L. *et al*., 2023) against a custom background consisting of our full list of 2735 proteins identified by DIA LC-MSMS. The full list of terms is supplied in **Supplemental file 21**. GO analysis was also performed to characterize over-representation of species enriched in the proximity of DCX(WT) relative to R178L, against the full 2735-protein background of our entire dataset, using g:profiler. Both analyses were performed using the BH correction and a standard significance threshold of p-value<0.05.

### Cell fixation and immunofluorescent staining

Cells were seeded at a density of ∼100,000 cells into 14 mm wells of poly-L-ornithine/laminin-coated #1.5 glass-bottom 35 mm dishes (MatTek) containing 200 μL of induction media. Following an initial 2-hour attachment window, dishes were flooded with induction media containing Y-27632 to support cell survival for the first 24-hours. Prior to fixing, cells were washed with warm DBPS without calcium or magnesium. Culture dishes were kept flat and never tilted during media changes or washes, to minimize tearing of neuronal processes. Neurons were fixed as previously described (Tint, I. *et al*., 2009), using freshly prepared 4% paraformaldehyde in PHEM buffer (60 mM PIPES, 25 mM HEPES, 10 mM EGTA, 2 mM MgCl2, pH 6.9 adjusted with KOH) to preserve cytoskeletal structures. Fixed cells were extracted with ice-cold methanol, quenched with 100 mM glycine in PBS, and blocked in 3% BSA containing 0.1% Triton X-100. Neurons were incubated with mouse anti-DCX (1:100; Santa Cruz) and either rabbit anti-MAP7D1 (1:50; Proteintech) or rabbit anti-EML4 (1:25; Cell Signaling Technologies). All samples were stained with anti-mouse Alexa Fluor 568 (Thermo Fisher Scientific), anti-rabbit Alexa Fluor 488 (Thermo Fisher Scientific), and TUBB3-647 (BioLegend), all at dilutions of 1:500. Samples were subsequently stained with DAPI (1:20,000), rinsed, and stored in PBS supplemented with 0.2% sodium azide.

### Widefield microscopy imaging

Images were acquired on a Axio Observer Z1 motorized inverted microscope (Zeiss) configured for Widefield Epifluorescence microscopy, controlled with Zen software (3.2 Blue edition, Zeiss), equipped with a 20x NA 0.8 Plan-Apochromat air objective and a 63x NA 1.4 Plan-Apochromat DIC oil objective (Zeiss). mages had a width of 2752 and a height of 2208 pixels. A set of images were acquired with the 20X air objective as z-stacks with 1 μm interval. The range of the z-stack varies depending on the cellular structures acquired. An equivalent set of images were acquired with the 63x oil objective. All 4 channels were excited with an X-Cite 120 LED light source (Excelitas) set to a light intensity specific to the channel, and their images were acquired on a Axiocam506 CCD camera (Zeiss) with channel-specific exposure times and 1×1 binning. DAPI (10% LED excitation; 150.00 ms exposure) wavelength selection was carried out with a BP 335/83 excitation filter (Zeiss), a FT 395 dichroic mirror (Zeiss) and a LP420 emission filter (Zeiss). The remaining channels were imaged as follows: 1) Alexa Fluor 488 – wavelength selection with a BP 455/95 excitation filter (Zeiss), FT 500 dichroic mirror (Zeiss) and BP505/55 emission filter (30% LED excitation, 500.00 ms exposure) - 2) Alexa Fluor 568 – wavelength selection with a BP 540/52 excitation filter (Zeiss), FT 580 dichroic mirror (Zeiss) and LP590 emission filter (30% LED excitation, 500.00 ms exposure) - 3) Alexa Fluor 647 – wavelength selection with a BP 643/88 excitation filter (Zeiss), FT695 dichroic mirror (Zeiss) and BP 700/50 emission filter (30% LED excitation, 500.00 ms exposure). Preliminary images were acquired using high-resolution confocal microscopy performed on a Leica Stellaris Confocal Laser Scanning Microscope (Stellaris 5) equipped with four highly sensitive HyD S detectors and a 63× oil-immersion objective.

### Microscopy analysis

Widefield microscopy hyperstacks were imported into FIJI (ImageJ) for image processing and analysis. 20x fields were utilized for all analysis, to ensure representation of results. For each field of view, regions of interest (ROIs) were manually defined on the raw images to delineate neurite processes, growth cones, somata, branch junctions, and background regions for each fluorescence channel. For neurite line-profile analysis, only clearly resolved, non-overlapping processes were traced, and fields containing fewer than 10 analyzable neurites were excluded. Background signal was measured separately for each channel within each field and subtracted from all fluorescence measurements. To account for developmental changes in protein expression, compartmental fluorescence intensities were normalized to the average maximum soma intensity measured for each channel within the corresponding field of view, as somata consistently exhibited the highest fluorescence intensity. Line-profile distances were normalized from 0–1 and fluorescence profiles were smoothed using a Savitzky– Golay filter (9-point window, second-order polynomial) to reduce high-frequency sampling noise while preserving profile morphology and peak position.

For MAP:TUBB3 analyses, matched MAP and TUBB3 line profiles were interpolated onto a common normalized distance axis, and pointwise MAP:TUBB3 intensity ratios were calculated along each neurite. Positions with normalized TUBB3 intensity below 0.02 were excluded to avoid unstable ratios arising from near-zero denominator values. The median ratio across all valid positions was used to represent the relative MAP:TUBB3 signal for each neurite. Pearson correlation coefficients were calculated from the unsmoothed, soma-normalized intensity profiles by comparing matched fluorescence values at corresponding positions along each neurite, providing a measure of the spatial correspondence between protein distributions.

## Key resources table

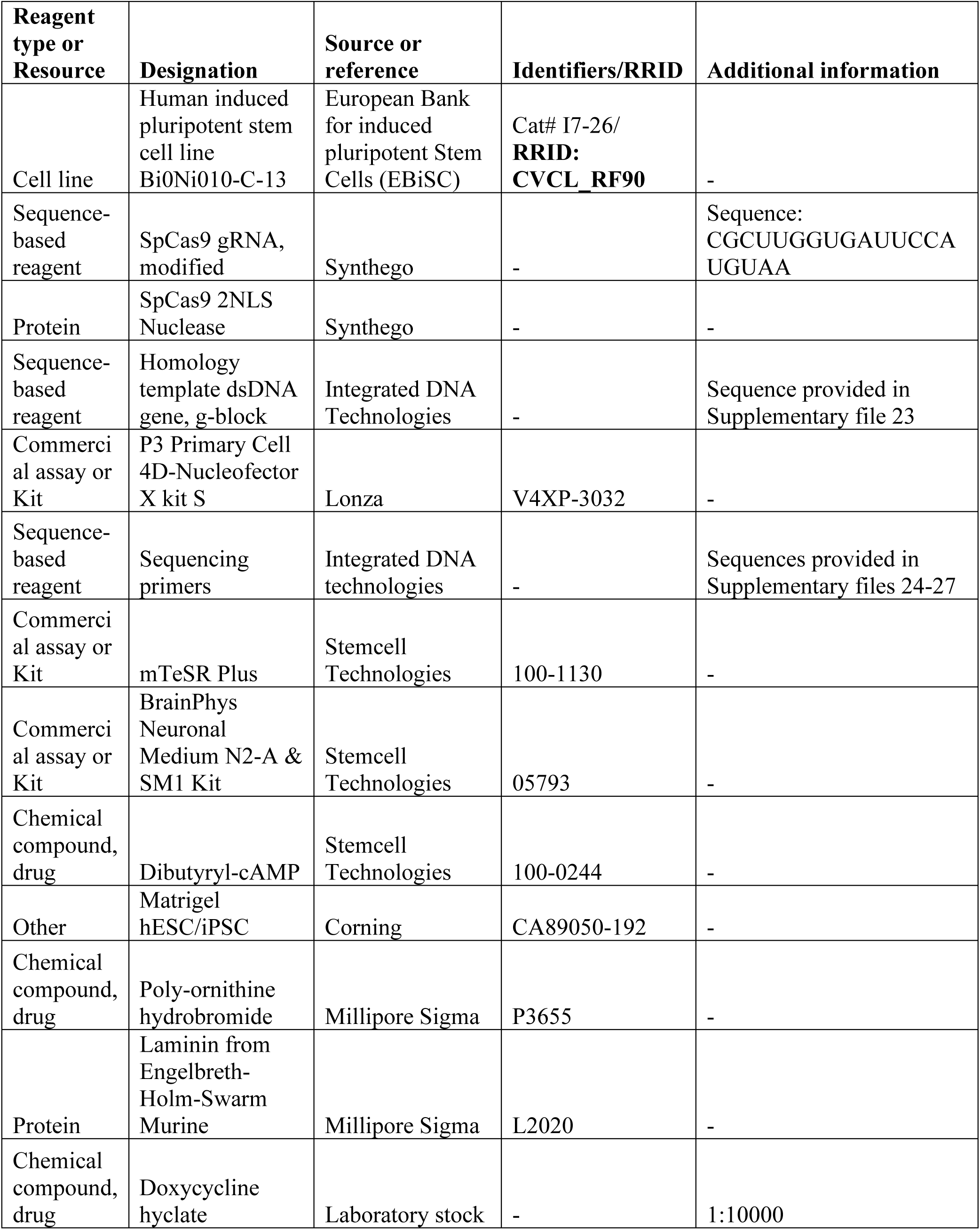
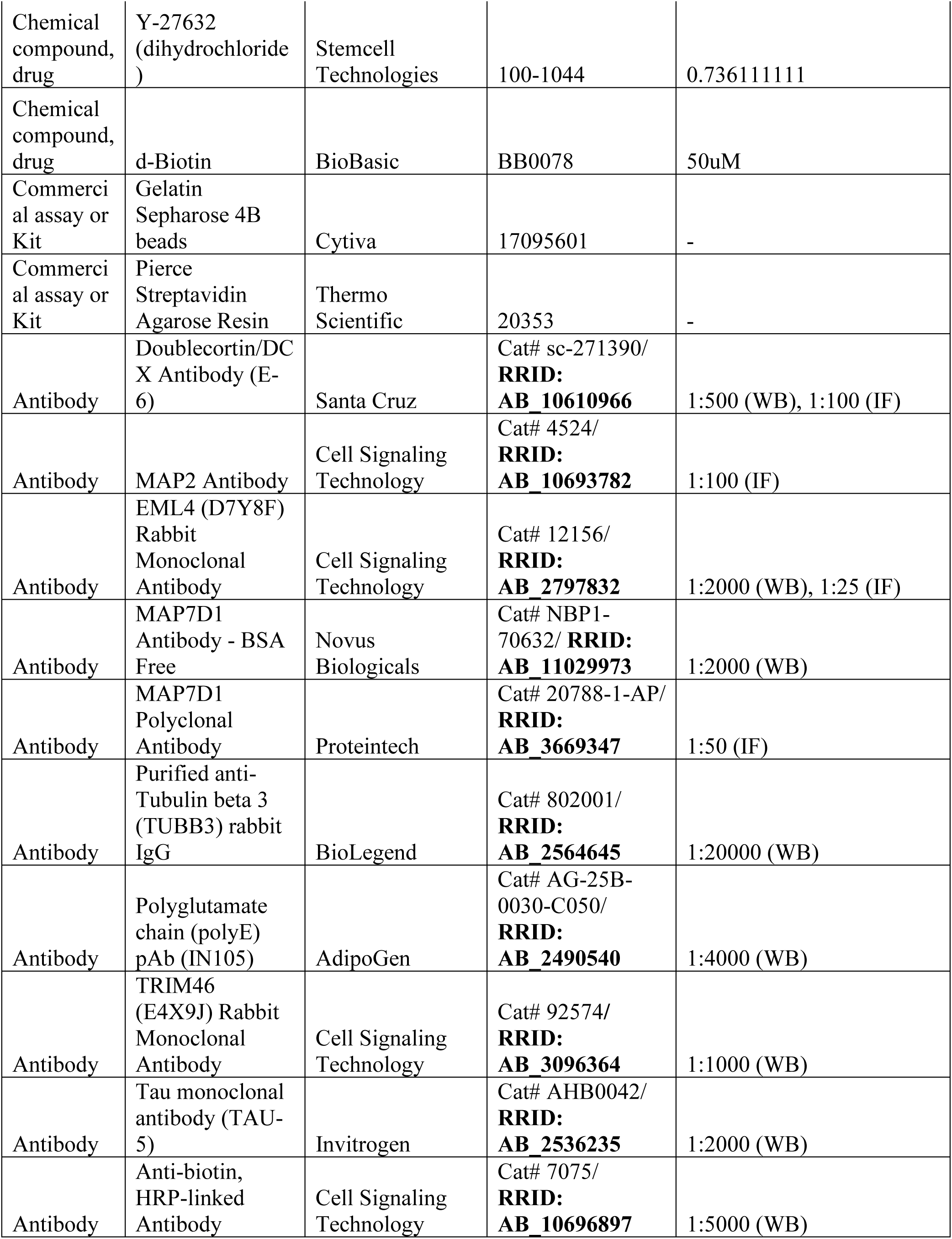
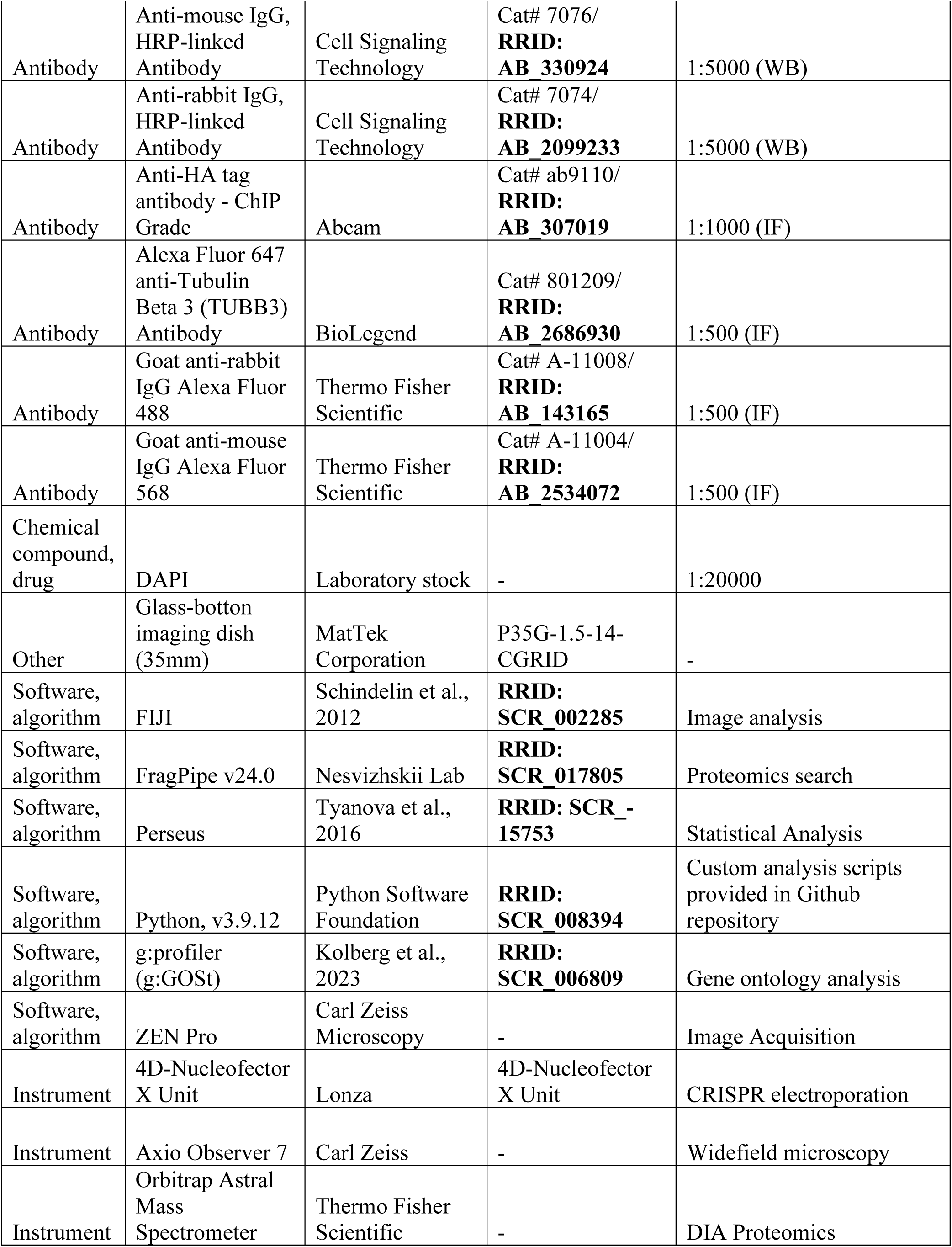

## Acknowledgements

We thank Kurt Dejgaard, Dr. rer. nat., and the Proteomics Core Facility at McGill University for their expertise and acquisition of preliminary data. We also thank Amy Wong, Jenna Cleyle, Laleh Mehboodi, and Lorne Taylor at the Proteomics and Molecular Analysis Platform (RI-MUHC) for technical support, including LC-MS sample preparation and data acquisition. Images were acquired through the McGill University Advanced Bioimaging Facility (ABIF; RRID: SCR_017697). We are indebted to the Centre de Recherche en Biologie Structurale (CRBS) and members of the CRBS community, who generously provided reagents, expertise, access to biophysical instrumentation, and financial support throughout this work. We thank Dr. H. Antonicka and members of the Shoubridge laboratory for providing the miniTurboID sequence map and their expertise in BioID. Finally, we thank lab members of J.-F. Trempe, L. Munter, A. Hendricks, S. Bechstedt, and G. Brouhard for their invaluable discussions, technical assistance, and contributions to experimental design. We are especially grateful to J.-F. Trempe for providing supervision, resources, and scientific guidance towards the completion of this project.

## Author contributions

B.L.K., L.M.M and J.-F.T. contributed to experimental design; B.L.K. conceived the project; G.J.B., L.M.M. and J.-F.T. provided supervisory support. B.L.K. and J.A.G. performed experiments and analysis. B.L.K. constructed analysis scripts. B.L.K. wrote the manuscript, with equal opportunity afforded to all authors for editing.

## Data availability

Mass spectrometry proteomics data have been deposited to the ProteomeXchange Consortium via the PRIDE partner repository (Perez-Riverol, Y. *et al*., 2025) with the dataset identifier PXD083262. Custom analysis and plotting scripts, including Jupyter notebooks used to generate figures, are publicly available in the DCX-proximity GitHub repository. Additional raw and source data files are available in the manuscript-associated Zenodo [Data repository](https://zenodo.org/records/22285498?preview=1&token=eyJhbGciOiJIUzUxMiJ9.eyJpZCI6IjJiM2JmMmJhLTMyNTctNGViMi04MDdjLWFmZDIwZmNkODYxZCIsImRhdGEiOnt9LCJyYW5kb20iOiI4MjI5MTk5NDk0ZDY4OTZjOTU0MGVhZGEyNDQwNmE0OCJ9.e2ZmKlgNONblp0fWDZ5Ko1E7cV5Fnmeo4k09kMKWsxdP5arcAh_ApIOsiaJMGbt08KgLVZ9QtNsI4hZZsDgDxQ).

## Bibliography

Adam, S. A. (2017). The Nucleoskeleton. Cold Spring Harbor Perspectives in Biology, 9(2). doi:10.1101/cshperspect.a023556

Antonicka, H., Lin, Z.-Y., Janer, A., Aaltonen, M. J., Weraarpachai, W., Gingras, A.-C., & Shoubridge, E. A. (2020). A High-Density Human Mitochondrial Proximity Interaction Network. Cell Metabolism, 32(3), 479–497.e479. doi:10.1016/j.cmet.2020.07.017

Arpag, G., Lawrence, E. J., Farmer, V. J., Hall, S. L., & Zanic, M. (2020). Collective effects of XMAP215, EB1, CLASP2, and MCAK lead to robust microtubule treadmilling. Proc Natl Acad Sci U S A, 117(23), 12847–12855. doi:10.1073/pnas.2003191117

Barnes, A. P., & Polleux, F. (2009). Establishment of Axon-Dendrite Polarity in Developing Neurons. Annual Review of Neuroscience, 32(Volume 32, 2009), 347–381. 10.1146/annurev.neuro.31.060407.125536

Bechstedt, S., & Brouhard, Gary J. (2012). Doublecortin Recognizes the 13-Protofilament Microtubule Cooperatively and Tracks Microtubule Ends. Developmental Cell, 23(1), 181–192. doi:10.1016/j.devcel.2012.05.006

Bechstedt, S., Lu, K., & Brouhard, G. J. (2014). Doublecortin Recognizes the Longitudinal Curvature of the Microtubule End and Lattice. Current Biology, 24(20), 2366–2375. doi:10.1016/j.cub.2014.08.039

Benjamini, Y., & Hochberg, Y. (1995). Controlling the False Discovery Rate: A Practical and Powerful Approach to Multiple Testing. Journal of the Royal Statistical Society: Series B (Methodological), 57(1), 289–300. 10.1111/j.2517-6161.1995.tb02031.x

Berger, S. L., Leo-Macias, A., Yuen, S., Khatri, L., Pfennig, S., Zhang, Y., Agullo-Pascual, E., Caillol, G., Zhu, M.-S., Rothenberg, E., Melendez-Vasquez, C. V., Delmar, M., Leterrier, C., & Salzer, J. L. (2018). Localized Myosin II Activity Regulates Assembly and Plasticity of the Axon Initial Segment. Neuron, 97(3), 555–570.e556. 10.1016/j.neuron.2017.12.039

Bielas, S. L., Serneo, F. F., Chechlacz, M., Deerinck, T. J., Perkins, G. A., Allen, P. B., Ellisman, M. H., & Gleeson, J. G. (2007). Spinophilin Facilitates Dephosphorylation of Doublecortin by PP1 to Mediate Microtubule Bundling at the Axonal Wrist. Cell, 129(3), 579–591. doi:10.1016/j.cell.2007.03.023

Bieling, P., Laan, L., Schek, H., Munteanu, E. L., Sandblad, L., Dogterom, M., Brunner, D., & Surrey, T. (2007). Reconstitution of a microtubule plus-end tracking system in vitro. Nature, 450(7172), 1100–1105. doi:10.1038/nature06386

Bott, C. J., McMahon, L. P., Keil, J. M., Choo Yap, C., Kwan, K. Y., & Winckler, B. (2020). Nestin Selectively Facilitates the Phosphorylation of the Lissencephaly-Linked Protein Doublecortin (DCX) by cdk5/p35 to Regulate Growth Cone Morphology and Sema3a Sensitivity in Developing Neurons. Journal of Neuroscience, 40(19), 3720–3740. doi:10.1523/JNEUROSCI.2471-19.2020

Branon, T. C., Bosch, J. A., Sanchez, A. D., Udeshi, N. D., Svinkina, T., Carr, S. A., Feldman, J. L., Perrimon, N., & Ting, A. Y. (2018). Efficient proximity labeling in living cells and organisms with TurboID. Nature Biotechnology, 36(9), 880–887. doi:10.1038/nbt.4201

Burger, D., Stihle, M., Sharma, A., Di Lello, P., Benz, J., D’Arcy, B., Debulpaep, M., Fry, D., Huber, W., Kremer, T., Laeremans, T., Matile, H., Ross, A., Rufer, A. C., Schoch, G., Steinmetz, M. O., Steyaert, J., Rudolph, M. G., Thoma, R., & Ruf, A. (2016). Crystal Structures of the Human Doublecortin C- and N-terminal Domains in Complex with Specific Antibodies. Journal of Biological Chemistry, 291(31), 16292–16306. doi:10.1074/jbc.M116.726547

Burnette, D. T., Ji, L., Schaefer, A. W., Medeiros, N. A., Danuser, G., & Forscher, P. (2008). Myosin II Activity Facilitates Microtubule Bundling in the Neuronal Growth Cone Neck. Developmental Cell, 15(1), 163–169. 10.1016/j.devcel.2008.05.016

Caceres, A., Banker, G., Steward, O., Binder, L., & Payne, M. (1984). MAP2 is localized to the dendrites of hippocampal neurons which develop in culture. Developmental Brain Research, 13(2), 314–318. 10.1016/0165-3806(84)90167-6

Cho, K. F., Branon, T. C., Udeshi, N. D., Myers, S. A., Carr, S. A., & Ting, A. Y. (2020). Proximity labeling in mammalian cells with TurboID and split-TurboID. Nature Protocols, 15(12), 3971–3999. doi:10.1038/s41596-020-0399-0

Cierpicki, T., Kim, M. H., Cooper, D. R., Derewenda, U., Bushweller, J. H., & Derewenda, Z. S. (2006). The DC-module of doublecortin: Dynamics, domain boundaries, and functional implications. *Proteins: Structure*, Function, and Bioinformatics, 64(4), 874–882. doi:10.1002/prot.21068

Combes, F., Loux, V., & Vandenbrouck, Y. (2021). GO Enrichment AnalysisGO enrichment analysis for Differential Proteomics Using ProteoREProteoRE. In D. Cecconi (Ed.), Proteomics Data Analysis (pp. 179–196). New York, NY: Springer US.

Dehmelt, L., & Halpain, S. (2004). The MAP2/Tau family of microtubule-associated proteins. Genome Biology, 6(1), 204. doi:10.1186/gb-2004-6-1-204

Dema, A., Charafeddine, R., Rahgozar, S., van Haren, J., & Wittmann, T. (2023). Growth cone advance requires EB1 as revealed by genomic replacement with a light-sensitive variant. Elife, 12, e84143. doi:10.7554/eLife.84143

Dema, A., Charafeddine, R. A., van Haren, J., Rahgozar, S., Viola, G., Jacobs, K. A., Kutys, M. L., & Wittmann, T. (2024). Doublecortin reinforces microtubules to promote growth cone advance in soft environments. Current Biology, 34(24), 5822–5832.e5825. doi:10.1016/j.cub.2024.10.063

Des Portes, V., Francis, F., Pinard, J. M., Desguerre, I., Moutard, M. L., Snoeck, I., Meiners, L. C., Capron, F., Cusmai, R., Ricci, S., Motte, J., Echenne, B., Ponsot, G., Dulac, O., Chelly, J., & Beldjord, C. (1998a). Doublecortin is the major gene causing X-linked subcortical laminar heterotopia (SCLH). Human Molecular Genetics, 7(7), 1063–1070. doi:10.1093/hmg/7.7.1063

Des Portes, V., Pinard, J. M., Billuart, P., Vinet, M. C., Koulakoff, A., Carrié, A., Gelot, A., Dupuis, E., Motte, J., Berwald-Netter, Y., Catala, M., Kahn, A., Beldjord, C., & Chelly, J. (1998b). A novel CNS gene required for neuronal migration and involved in X-linked subcortical laminar heterotopia and lissencephaly syndrome. Cell, 92(1), 51–61. doi:10.1016/S0092-8674(00)80898-3

Fu, X., Brown, K. J., Yap, C. C., Winckler, B., Jaiswal, J. K., & Liu, J. S. (2013). Doublecortin (Dcx) family proteins regulate filamentous actin structure in developing neurons. Journal of Neuroscience, 33(2), 709–721. doi:10.1523/JNEUROSCI.4603-12.2013

Fu, X., Rao, L., Li, P., Liu, X., Wang, Q., Son, A. I., Gennerich, A., & Liu, J. S.-H. (2022). Doublecortin and JIP3 are neural-specific counteracting regulators of dynein-mediated retrograde trafficking. Elife, 11, e82218. doi:10.7554/eLife.82218

Gavoci, A., Zhiti, A., Rusková, M., Magiera, M. M., Wang, M., Ziegler, K. A., Hausrat, T. J., Ugwuja, A. I., Chakraborty, S., Engelhardt, S., Kneussel, M., Balastik, M., Janke, C., Misgeld, T., & Brill, M. S. (2025). Polyglutamylation of microtubules drives neuronal remodeling. Nature Communications, 16(1), 5384. doi:10.1038/s41467-025-60855-6

Gleeson, J. G., Allen, K. M., Fox, J. W., Lamperti, E. D., Berkovic, S., Scheffer, I., Cooper, E. C., Dobyns, W. B., Minnerath, S. R., Ross, M. E., & Walsh, C. A. (1998). doublecortin, a Brain-Specific Gene Mutated in Human X-Linked Lissencephaly and Double Cortex Syndrome, Encodes a Putative Signaling Protein. Cell, 92(1), 63–72. doi:10.1016/S0092-8674(00)80899-5

Gleeson, J. G., Lin, P. T., Flanagan, L. A., & Walsh, C. A. (1999a). Doublecortin Is a Microtubule-Associated Protein and Is Expressed Widely by Migrating Neurons. Neuron, 23(2), 257–271. doi:10.1016/S0896-6273(00)80778-3

Gleeson, J. G., Minnerath, S. R., Fox, J. W., Allen, K. M., Luo, R. F., Hong, S. E., Berg, M. J., Kuzniecky, R., Reitnauer, P. J., Borgatti, R., Mira, A. P., Guerrini, R., Holmes, G. L., Rooney, C. M., Berkovic, S., Scheffer, I., Cooper, E. C., Ricci, S., Cusmai, R., Crawford, T. O., Leroy, R., Andermann, E., Wheless, J. W., Dobyns, W. B., Walsh, C. A., &, et al. (1999b). Characterization of mutations in the gene doublecortin in patients with double cortex syndrome. Ann Neurol, 45(2), 146–153. doi:10.1002/1531-8249(199902)45:2<146::aid-ana3>3.0.co;2-n

Hindley, C., Ali, F., McDowell, G., Cheng, K., Jones, A., Guillemot, F., & Philpott, A. (2012). Post-translational modification of Ngn2 differentially affects transcription of distinct targets to regulate the balance between progenitor maintenance and differentiation. Development, 139(10), 1718–1723. doi:10.1242/dev.077552

Hooikaas, P. J., Martin, M., Mühlethaler, T., Kuijntjes, G.-J., Peeters, C. A. E., Katrukha, E. A., Ferrari, L., Stucchi, R., Verhagen, D. G. F., van Riel, W. E., Grigoriev, I., Altelaar, A. F. M., Hoogenraad, C. C., Rüdiger, S. G. D., Steinmetz, M. O., Kapitein, L. C., & Akhmanova, A. (2019). MAP7 family proteins regulate kinesin-1 recruitment and activation. Journal of Cell Biology, 218(4), 1298–1318. doi:10.1083/jcb.201808065

Houtman, S. H., Rutteman, M., De Zeeuw, C. I., & French, P. J. (2007). Echinoderm microtubule-associated protein like protein 4, a member of the echinoderm microtubule-associated protein family, stabilizes microtubules. Neuroscience, 144(4), 1373–1382. 10.1016/j.neuroscience.2006.11.015

Hulme, A. J., Maksour, S., St-Clair Glover, M., Miellet, S., & Dottori, M. (2022). Making neurons, made easy: The use of Neurogenin-2 in neuronal differentiation. Stem Cell Reports, 17(1), 14–34. 10.1016/j.stemcr.2021.11.015

Irgen-Gioro, S., Yoshida, S., Walling, V., & Chong, S. (2022). Fixation can change the appearance of phase separation in living cells. Elife, 11, e79903. doi:10.7554/eLife.79903

Janke, C., & Kneussel, M. (2010). Tubulin post-translational modifications: encoding functions on the neuronal microtubule cytoskeleton. Trends in Neurosciences, 33(8), 362–372. doi:10.1016/j.tins.2010.05.001

Jin, J., Suzuki, H., Hirai, S. I., Mikoshiba, K., & Ohshima, T. (2010). JNK phosphorylates Ser332 of doublecortin and regulates its function in neurite extension and neuronal migration. Developmental Neurobiology, 70(14), 929–942. doi:10.1002/dneu.20833

Kappeler, C., Saillour, Y., Baudoin, J. P., Tuy, F. P. D., Alvarez, C., Houbron, C., Gaspar, P., Hamard, G., Chelly, J., Métin, C., & Francis, F. (2006). Branching and nucleokinesis defects in migrating interneurons derived from doublecortin knockout mice. Human Molecular Genetics, 15(9), 1387–1400. doi:10.1093/hmg/ddl062

Kevenaar, J. T., & Hoogenraad, C. C. (2015). The axonal cytoskeleton: from organization to function. Frontiers in Molecular Neuroscience, Volume 8 - 2015. doi:10.3389/fnmol.2015.00044

Kikuchi, K., Sakamoto, Y., Uezu, A., Yamamoto, H., Ishiguro, K.-i., Shimamura, K., Saito, T., Hisanaga, S.-i., & Nakanishi, H. (2022). Map7D2 and Map7D1 facilitate microtubule stabilization through distinct mechanisms in neuronal cells. Life Science Alliance, 5(8), e202201390. doi:10.26508/lsa.202201390

Kim, M. H., Cierpicki, T., Derewenda, U., Krowarsch, D., Feng, Y., Devedjiev, Y., Dauter, Z., Walsh, C. A., Otlewski, J., Bushweller, J. H., & Derewenda, Z. S. (2003). The DCX-domain tandems of doublecortin and doublecortin-like kinase. Nature Structural & Molecular Biology, 10(5), 324–333. doi:10.1038/nsb918

Koizumi, H., Fujioka, H., Togashi, K., Thompson, J., Yates III, J. R., Gleeson, J. G., & Emoto, K. (2017). DCLK1 phosphorylates the microtubule-associated protein MAP7D1 to promote axon elongation in cortical neurons. Developmental Neurobiology, 77(4), 493–510. 10.1002/dneu.22428

Koizumi, H., Higginbotham, H., Poon, T., Tanaka, T., Brinkman, B. C., & Gleeson, J. G. (2006). Doublecortin maintains bipolar shape and nuclear translocation during migration in the adult forebrain. Nat Neurosci, 9(6), 779–786. doi:10.1038/nn1704

Kolberg, L., Raudvere, U., Kuzmin, I., Adler, P., Vilo, J., & Peterson, H. (2023). g:Profiler—interoperable web service for functional enrichment analysis and gene identifier mapping (2023 update). Nucleic Acids Research, 51(W1), W207–W212. doi:10.1093/nar/gkad347

Kuo, Y.-W., & Howard, J. (2021). Cutting, Amplifying, and Aligning Microtubules with Severing Enzymes. Trends in Cell Biology, 31(1), 50–61. doi:10.1016/j.tcb.2020.10.004

Li, P., Li, L., Yu, B., Wang, X., Wang, Q., Lin, J., Zheng, Y., Zhu, J., He, M., Xia, Z., Tu, M., Liu, J. S., Lin, Z., & Fu, X. (2021). Doublecortin facilitates the elongation of the somatic Golgi apparatus into proximal dendrites. Molecular Biology of the Cell, 32(5), 422–434. doi:10.1091/mbc.E19-09-0530

Liao, Y.-C., Tsai, M.-H., Chao, N.-H., Chang, Y.-S., Lin, T.-W., Lin, I. H., Hou, P.-S., Wang, W.-J., & Tsai, J.-W. (2026). CEP170 as a novel molecular link between centrosomal function and cerebral cortical development. Journal of Biomedical Science, 33(1), 33. doi:10.1186/s12929-026-01236-z

Liu, Judy S., Schubert, Christian R., Fu, X., Fourniol, Franck J., Jaiswal, Jyoti K., Houdusse, A., Stultz, Collin M., Moores, Carolyn A., & Walsh, Christopher A. (2012). Molecular Basis for Specific Regulation of Neuronal Kinesin-3 Motors by Doublecortin Family Proteins. Molecular Cell, 47(5), 707–721. doi:10.1016/j.molcel.2012.06.025

Locard-Paulet, M., Doncheva, N. T., Morris, J. H., & Jensen, L. J. (2024). Functional Analysis of MS-Based Proteomics Data: From Protein Groups to Networks. Molecular & Cellular Proteomics, 23(12). doi:10.1016/j.mcpro.2024.100871

Madgwick, A., Fort, P., Hanson, P. S., Thibault, P., Gaudreau, M.-C., Lutfalla, G., Möröy, T., Abou Elela, S., Chaudhry, B., Elliott, D. J., Morris, C. M., & Venables, J. P. (2015). Neural Differentiation Modulates the Vertebrate Brain Specific Splicing Program. PLOS ONE, 10(5), e0125998. doi:10.1371/journal.pone.0125998

Manka, S. W., & Moores, C. A. (2020). Pseudo-repeats in doublecortin make distinct mechanistic contributions to microtubule regulation. EMBO Rep, 21(12), e51534. doi:10.15252/embr.202051534

May, D. G., Scott, K. L., Campos, A. R., & Roux, K. J. (2020). Comparative Application of BioID and TurboID for Protein-Proximity Biotinylation. Cells, 9(5). doi:10.3390/cells9051070

McAlear, T. S., & Bechstedt, S. (2022). The mitotic spindle protein CKAP2 potently increases formation and stability of microtubules. Elife, 11, e72202. doi:10.7554/eLife.72202

Monroy, B. Y., Sawyer, D. L., Ackermann, B. E., Borden, M. M., Tan, T. C., & Ori-McKenney, K. M. (2018). Competition between microtubule-associated proteins directs motor transport. Nat Commun, 9(1), 1487. doi:10.1038/s41467-018-03909-2

Monroy, B. Y., Tan, T. C., Oclaman, J. M., Han, J. S., Simo, S., Niwa, S., Nowakowski, D. W., McKenney, R. J., & Ori-McKenney, K. M. (2020). A Combinatorial MAP Code Dictates Polarized Microtubule Transport. Dev Cell, 53(1), 60–72 e64. doi:10.1016/j.devcel.2020.01.029

Moores, C. A., Perderiset, M., Francis, F., Chelly, J., Houdusse, A., & Milligan, R. A. (2004). Mechanism of microtubule stabilization by doublecortin. Mol Cell, 14(6), 833–839. doi:10.1016/j.molcel.2004.06.009

Muñoz-Estrada, J., Mostafania, A., Halwatura, L., Haghani, A., Jiang, Y., & Meyer, J. G. (2026). Optimizing NGN2 Dosage Enhances the Neuronal Enrichment of iPSC-derived Neuronal Cultures. Molecular & Cellular Proteomics. doi:10.1016/j.mcpro.2026.101604

Perez-Riverol, Y., Bandla, C., Kundu, Deepti J., Kamatchinathan, S., Bai, J., Hewapathirana, S., John, Nithu S., Prakash, A., Walzer, M., Wang, S., & Vizcaíno, Juan A. (2025). The PRIDE database at 20 years: 2025 update. Nucleic Acids Research, 53(D1), D543–D553. doi:10.1093/nar/gkae1011

Poirier, K., Saillour, Y., Bahi-Buisson, N., Jaglin, X. H., Fallet-Bianco, C., Nabbout, R., Castelnau-Ptakhine, L., Roubertie, A., Attie-Bitach, T., Desguerre, I., Genevieve, D., Barnerias, C., Keren, B., Lebrun, N., Boddaert, N., Encha-Razavi, F., & Chelly, J. (2010). Mutations in the neuronal β-tubulin subunit TUBB3 result in malformation of cortical development and neuronal migration defects. Human Molecular Genetics, 19(22), 4462–4473. doi:10.1093/hmg/ddq377

Pollmann, M., Parwaresch, R., Adam-Klages, S., Kruse, M.-L., Buck, F., & Heidebrecht, H.-J. (2006). Human EML4, a novel member of the EMAP family, is essential for microtubule formation. Experimental Cell Research, 312(17), 3241–3251. 10.1016/j.yexcr.2006.06.035

Qiang, L., Yu, W., Andreadis, A., Luo, M., & Baas, P. W. (2006). Tau Protects Microtubules in the Axon from Severing by Katanin. The Journal of Neuroscience, 26(12), 3120–3129. doi:10.1523/jneurosci.5392-05.2006

Qu, C., Dwyer, T., Shao, Q., Yang, T., Huang, H., & Liu, G. (2013). Direct binding of TUBB3 with DCC couples netrin-1 signaling to intracellular microtubule dynamics in axon outgrowth and guidance. Journal of Cell Science, 126(14), 3070–3081. doi:10.1242/jcs.122184

Rafiei, A., Cruz Tetlalmatzi, S., Edrington, C. H., Lee, L., Crowder, D. A., Saltzberg, D. J., Sali, A., Brouhard, G., & Schriemer, D. C. (2022). Doublecortin engages the microtubule lattice through a cooperative binding mode involving its C-terminal domain. Elife, 11, e66975. doi:10.7554/eLife.66975

Rufer, A. C., Kusznir, E., Burger, D., Stihle, M., Ruf, A., & Rudolph, M. G. (2018). Domain swap in the C-terminal ubiquitin-like domain of human doublecortin. Acta Crystallogr D Struct Biol, 74(5), 450–462. doi:10.1107/S2059798318004813

Saadi, I., Goering, J. P., Hufft-Martinez, B. M., & Tran, P. V. (2023). SPECC1L: a cytoskeletal protein that regulates embryonic tissue dynamics. Biochemical Society Transactions, 51(3), 949–958. doi:10.1042/bst20220461

Sébastien, M., Paquette, A. L., Prowse, E. N. P., Hendricks, A. G., & Brouhard, G. J. (2025). Doublecortin restricts neuronal branching by regulating tubulin polyglutamylation. Nature Communications, 16(1), 1749. doi:10.1038/s41467-025-56951-2

Shao, Q., Yang, T., Huang, H., Majumder, T., Khot, B. A., Khouzani, M. M., Alarmanazi, F., Gore, Y. K., & Liu, G. (2019). Disease-associated mutations in human TUBB3 disturb netrin repulsive signaling. PLOS ONE, 14(6), e0218811. doi:10.1371/journal.pone.0218811

Shatery Nejad, N., Boczkowska, M., Hilal, R., Fregoso, F. E., Barrie, K. R., Rebowski, G., Saks, A. J., Gautreau, A. M., De La Cruz, E. M., & Dominguez, R. (2025). Mechanism of Arp2/3 complex branch disassembly by human Coro7. Nature Communications, 16(1), 9809. doi:10.1038/s41467-025-64787-z

Slepak, T. I., Salay, L. D., Lemmon, V. P., & Bixby, J. L. (2012). Dyrk kinases regulate phosphorylation of doublecortin, cytoskeletal organization, and neuronal morphology. Cytoskeleton, 69(7), 514–527. 10.1002/cm.21021

Stouffer, M. A., Khalaf-Nazzal, R., Cifuentes-Diaz, C., Albertini, G., Bandet, E., Grannec, G., Lavilla, V., Deleuze, J. F., Olaso, R., Nosten-Bertrand, M., & Francis, F. (2022). Doublecortin mutation leads to persistent defects in the Golgi apparatus and mitochondria in adult hippocampal pyramidal cells. Neurobiology of Disease, 168, 105702. doi:10.1016/j.nbd.2022.105702

Tanaka, T., Serneo, F. F., Higgins, C., Gambello, M. J., Wynshaw-Boris, A., & Gleeson, J. G. (2004). Lis1 and doublecortin function with dynein to mediate coupling of the nucleus to the centrosome in neuronal migration. J Cell Biol, 165(5), 709–721. doi:10.1083/jcb.200309025

Tint, I., Jean, D., Baas, P. W., & Black, M. M. (2009). Doublecortin associates with microtubules preferentially in regions of the axon displaying actin-rich protrusive structures. Journal of Neuroscience, 29(35), 10995–11010. doi:10.1523/JNEUROSCI.3399-09.2009

Toriyama, M., Mizuno, N., Fukami, T., Iguchi, T., Toriyama, M., Tago, K., & Itoh, H. (2012). Phosphorylation of doublecortin by protein kinase A orchestrates microtubule and actin dynamics to promote neuronal progenitor cell migration. Journal of Biological Chemistry, 287(16), 12691–12702. doi:10.1074/jbc.M111.316307

Tsukada, M., Prokscha, A., Ungewickell, E., & Eichele, G. (2005). Doublecortin Association with Actin Filaments Is Regulated by Neurabin II. Journal of Biological Chemistry, 280(12), 11361–11368. doi:10.1074/jbc.M405525200 10.1074/jbc.M405525200

Uversky, V. N. (2002). Natively unfolded proteins: A point where biology waits for physics. Protein Science, 11(4), 739–756. 10.1110/ps.4210102

Verhey, K. J., & Gaertig, J. (2007). The Tubulin Code. Cell Cycle, 6(17), 2152–2160. doi:10.4161/cc.6.17.4633

Vinopal, S., & Bradke, F. (2025). Centrosomal and acentrosomal microtubule nucleation during neuronal development. Current Opinion in Neurobiology, 92, 103016. 10.1016/j.conb.2025.103016

Vinopal, S., Dupraz, S., Alfadil, E., Pietralla, T., Bendre, S., Stiess, M., Falk, S., Camargo Ortega, G., Maghelli, N., Tolić, I. M., Smejkal, J., Götz, M., & Bradke, F. (2023). Centrosomal microtubule nucleation regulates radial migration of projection neurons independently of polarization in the developing brain. Neuron, 111(8), 1241–1263.e1216. doi:10.1016/j.neuron.2023.01.020

Wong, H. H.-W., Chou, C. Y. C., Watt, A. J., & Sjöström, P. J. (2023). Comparing mouse and human brains. Elife, 12, e90017. doi:10.7554/eLife.90017

Yap, C. C., Digilio, L., McMahon, L., Roszkowska, M., Bott, C. J., Kruczek, K., & Winckler, B. (2016). Different doublecortin (DCX) patient alleles show distinct phenotypes in cultured neurons: Evidence for divergent loss-of-function and "off-pathway" cellular mechanisms. Journal of Biological Chemistry, 291(52), 26613–26626. doi:10.1074/jbc.M116.760777

Yu, W., Qiang, L., Solowska, J. M., Karabay, A., Korulu, S., Baas, P. W., & Holzbaur, E. (2008). The Microtubule-severing Proteins Spastin and Katanin Participate Differently in the Formation of Axonal Branches. Molecular Biology of the Cell, 19(4), 1485–1498. doi:doi:10.1091/mbc.e07-09-0878

Zhang, X., Lei, K., Yuan, X., Wu, X., Zhuang, Y., Xu, T., Xu, R., & Han, M. (2009). SUN1/2 and Syne/Nesprin-1/2 Complexes Connect Centrosome to the Nucleus during Neurogenesis and Neuronal Migration in Mice. Neuron, 64(2), 173–187. doi:10.1016/j.neuron.2009.08.018

