## Supplemental File 1 (Supporting Figures) for "Doublecortin mutation disrupts lattice-dependent proximity to reveal cytoskeletal networks in developing neurons"

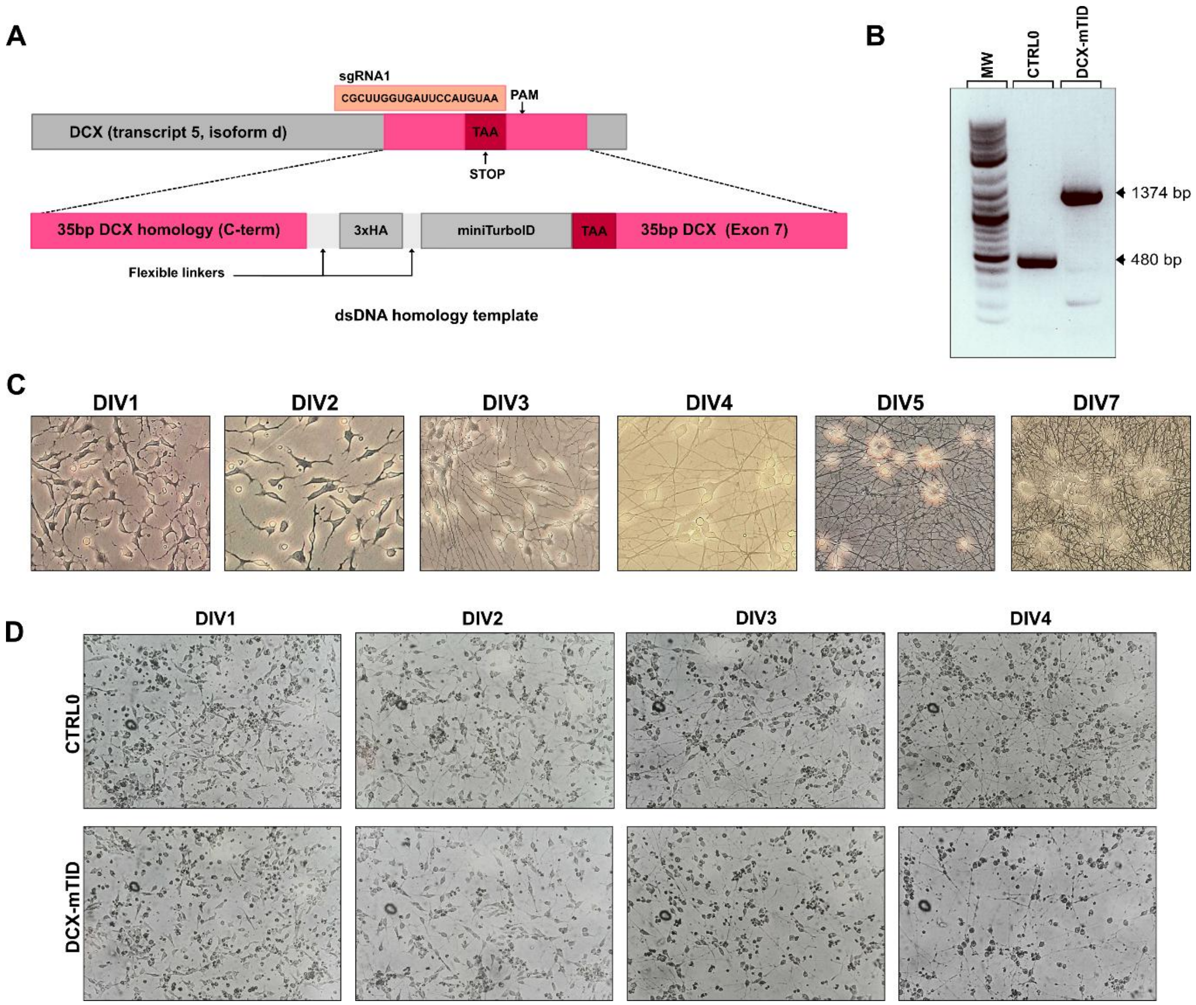


**Figure 1 – Supplement 1:** ***CRISPR-Cas9 knock-in and cell line validation.*** **(A)** Schematic illustrating knock-in strategy, using dsDNA homology regions flanking coding regions for HA tag and miniTurboID, with homology encoding the endogenous STOP codon for DCX transcript 5. **(B)** Agarose gel used for genotyping, showing the presence (DCX-mTID) and absence (CTRL0) of ~900bp, the length of the donor dsDNA. **(C)** Differentiation series of Brightfield images demonstrating morphology of neurons from induction (DIV0) to maturity (DIV7) in unedited CTRL0 cells. **(D)** Differentiation series of Polarized Light microscopy images comparing morphological stages between knock-in (DCX-mTID) and unedited (CTRL0) cells.

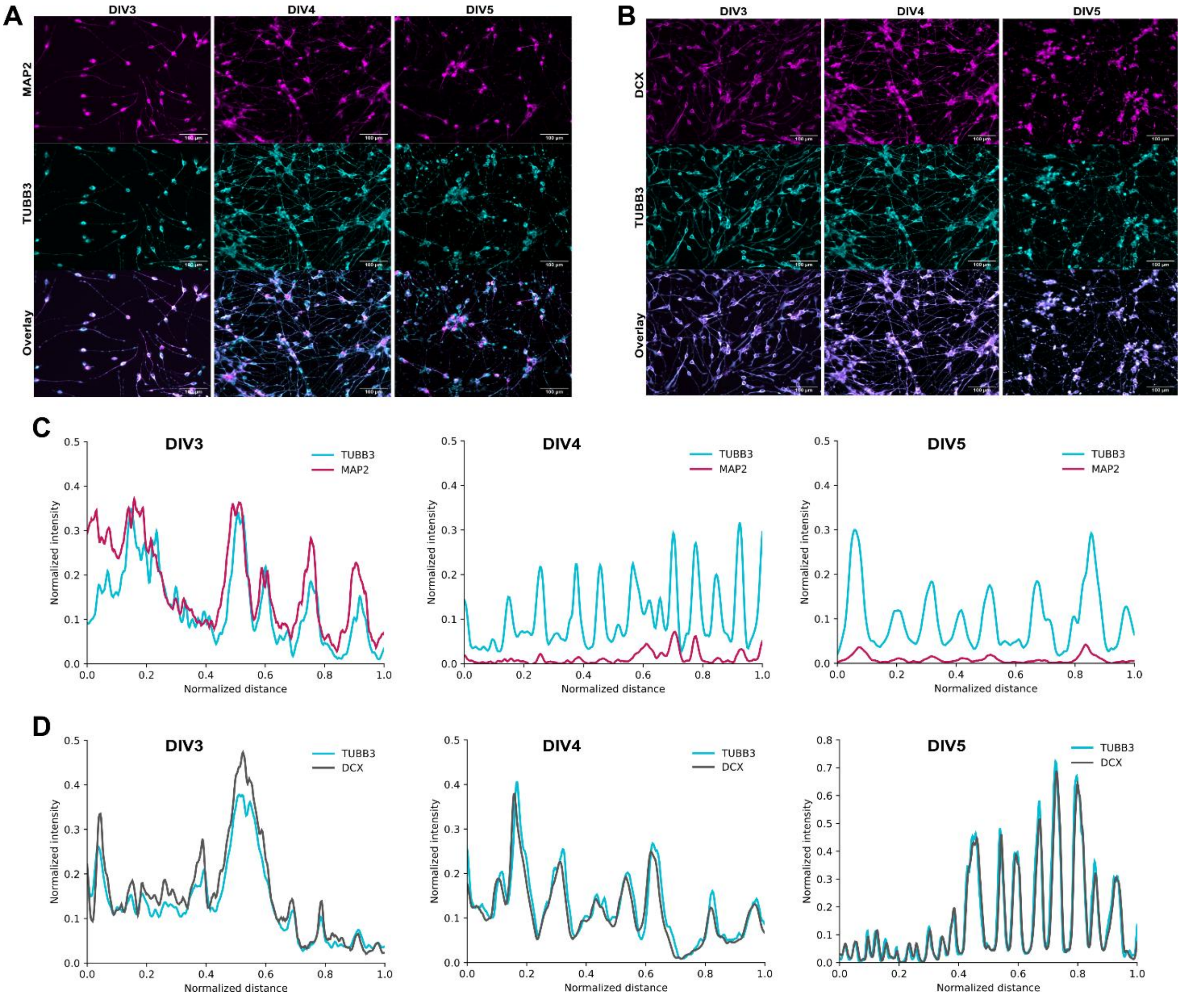


**Figure 1 – Supplement 2:** ***Validating DCX localization throughout differentiation.*** **(A)** Widefield fluorescence 20x images showing the MAP2 (magenta) localization overlaid with TUBB3 (cyan), represented by neurite profiles in (C). **(B)** Widefield 20x images showing DCX (magenta) localization overlaid with TUBB3 (cyan), represented by neurite profiles in (D). **(C-D)** Representative line profiles showing the normalized intensity of MAP2 (magenta) (C) or DCX (grey) (D) measured along neurites, overlaid with measured intensity for TUBB3 (cyan) at each DIV. Line profiles are representative of a population of >10 neurites per field, >3 fields per biological replicate, and >2 biological replicates.

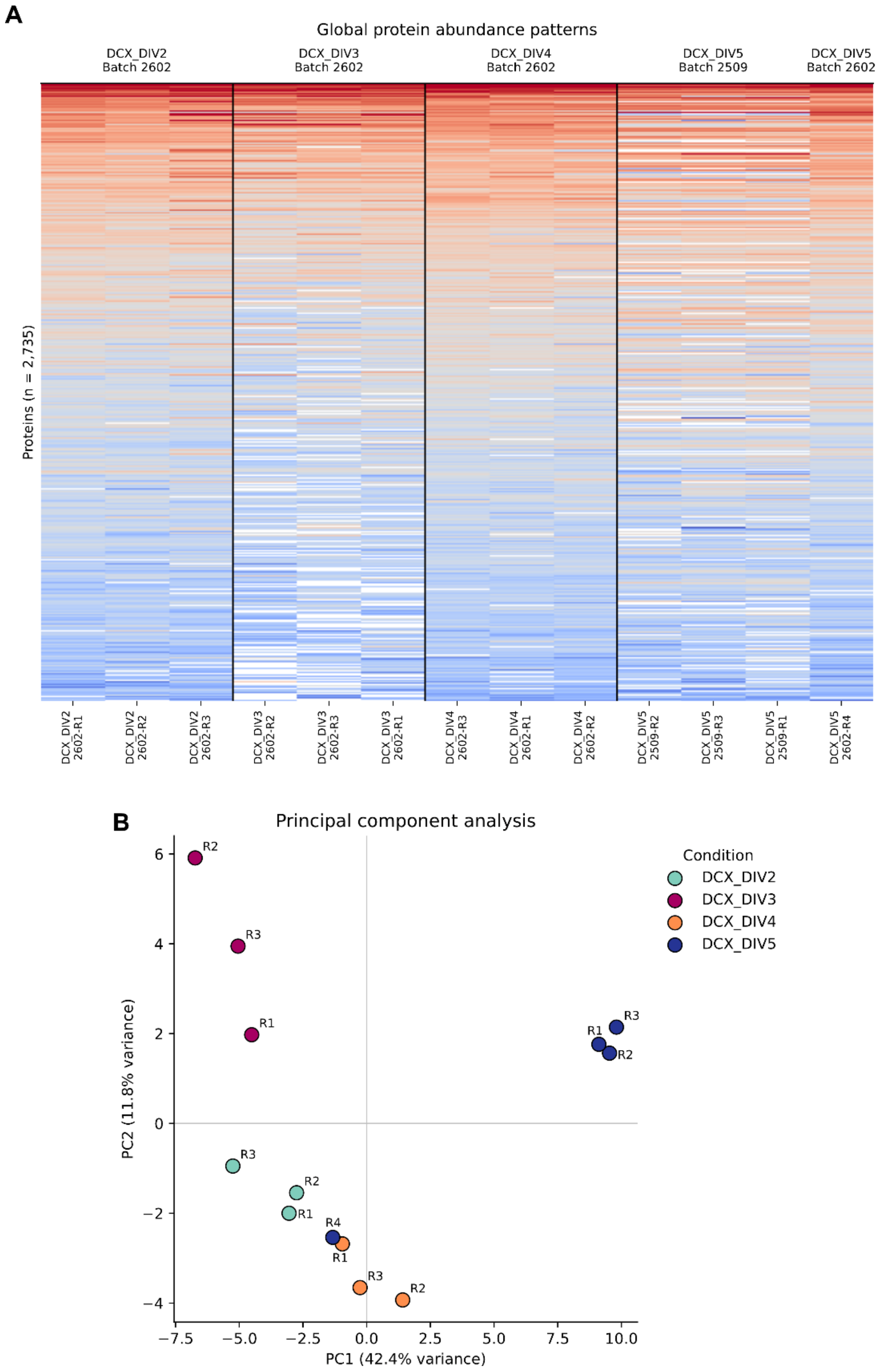


**Figure 2 – Supplement 1:** ***Batch-level and replicate behavior of the bulk proteome.*** **(A)** Heat map showing hierarchical clustering of protein intensities across individual samples. Biological replicates (R1–R5) and MS acquisition batches (2509 and 2602) are indicated for DIV2 (n = 3), DIV3 (n = 3), DIV4 (n = 3), and DIV5 WT-DCX (n = 4). **(B)** Principal component analysis (PCA) showing clustering of biological replicates by developmental stage and genotype.

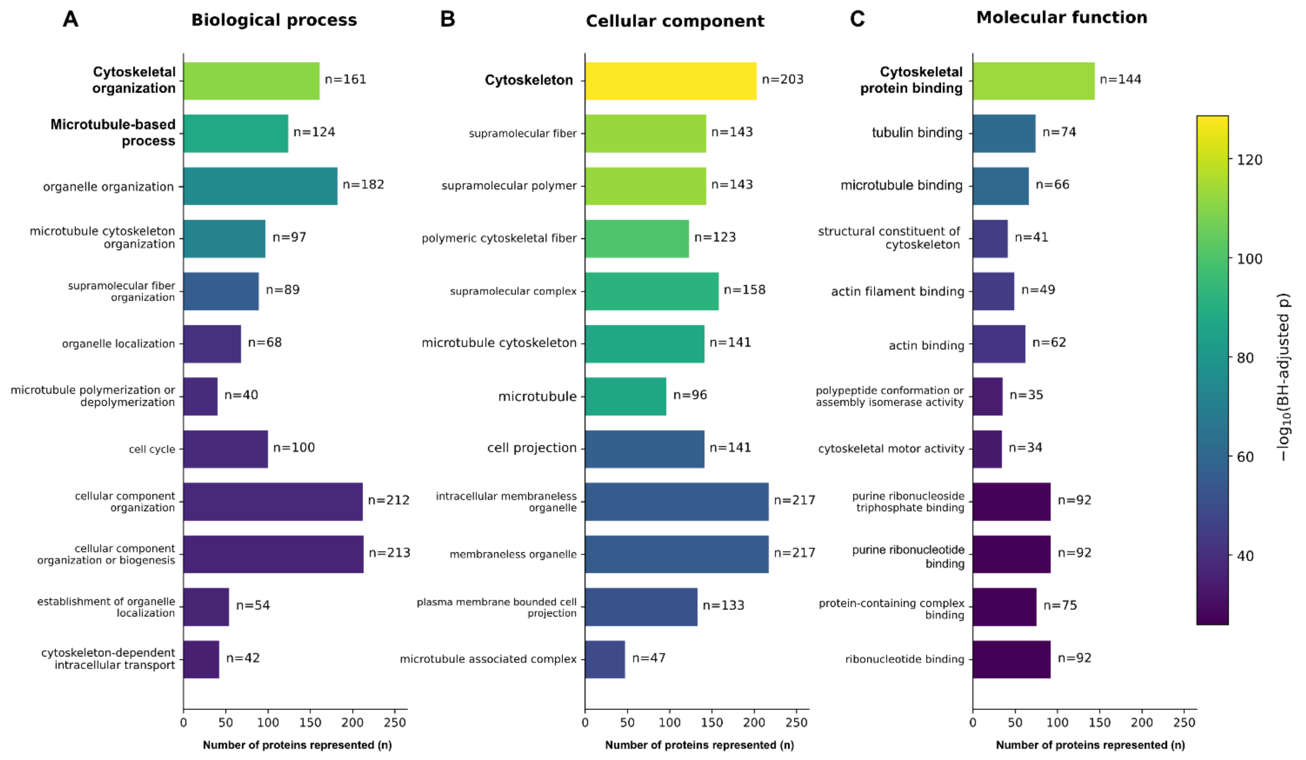


**Figure 2 – Supplement 2:** ***Gene Ontology over-representation analysis (GO-ORA) of the curated Cytoskeletal database.*** **(A-C)** GO enrichment plot showing the 12 most-enriched terms according to Biological Process (BP; panel A), Cellular Component (CC; panel B), and Molecular Function (MF; panel C) categories. Bar length represents the number of proteins from the curated cytoskeletal subset annotated to each GO term (n); Bar colour represents enrichment significance, expressed as $-\log_{10}$ of the Benjamini-Hochberg-adjusted *p*-value. Input list included proteins in manually curated Cytoskeletal database against all identified proteins in DIV1-5 dataset. Full list of terms is detailed in **Supplemental File 21**.

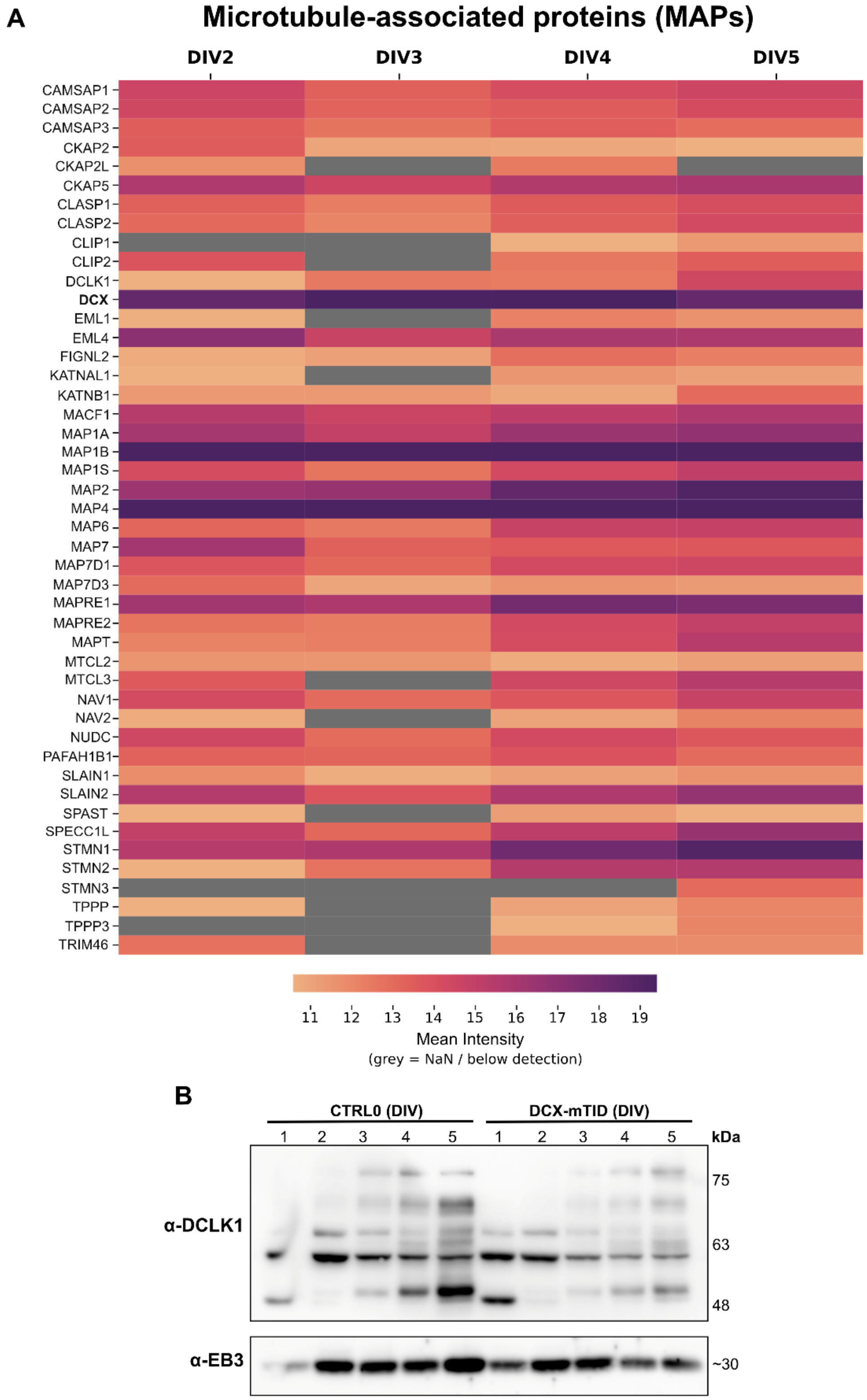


**Figure 3 – Supplement 1:** ***Cytoskeletal Atlas I – MAPs.*** **A)** Protein-level intensity heatmap for proteins compiled in Cytoskeleton database, as catalogued in Supplemental File 3 (Perseus) and Supplemental File 18. **(B)** Immunoblot evidence in neuronal lysates for select MAPs: doublecortin-like kinase 1 (DCLK1), end-binding protein EB3/MAPRE3.

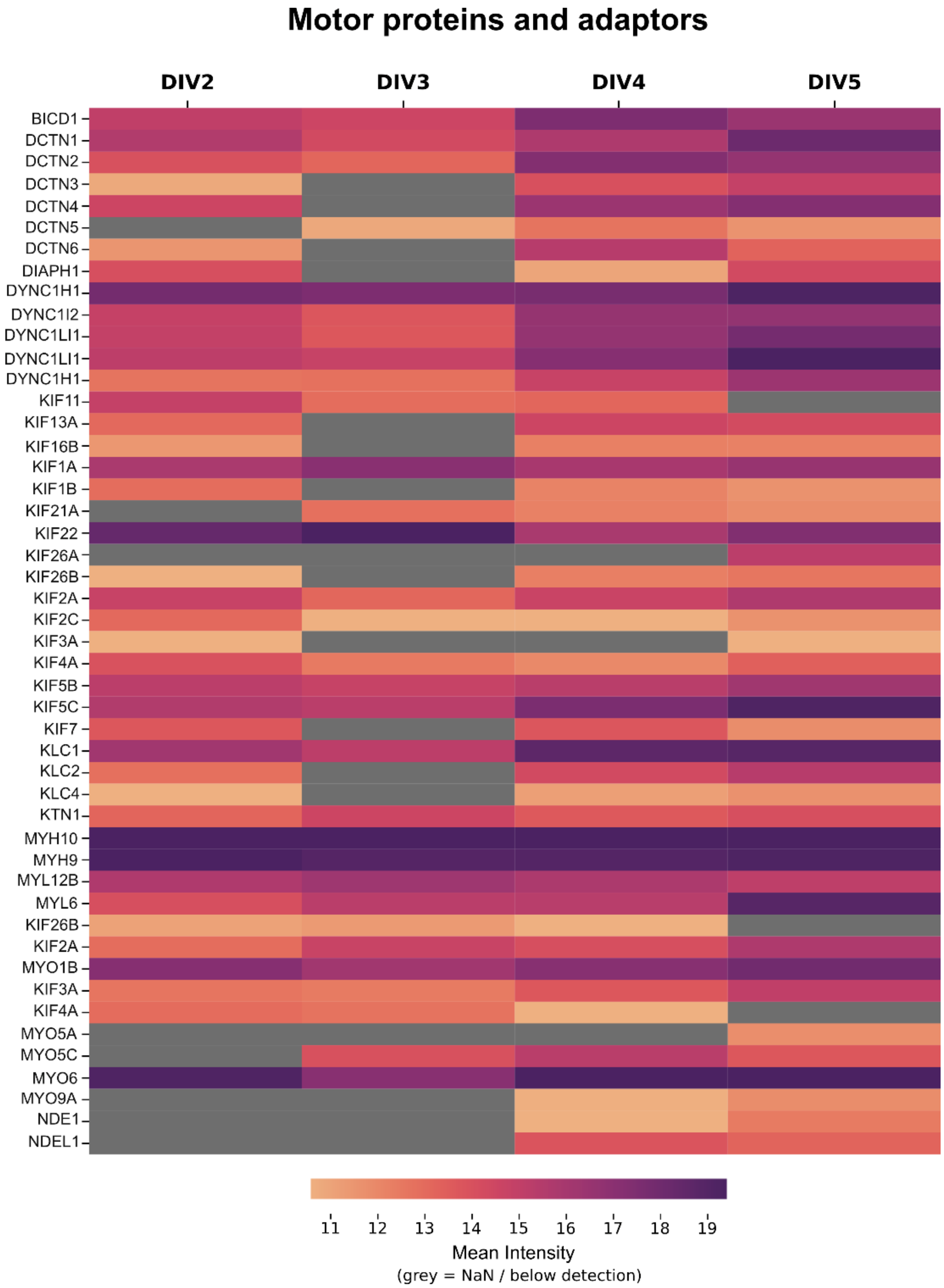


**Figure 3 – Supplement 2:** ***Cytoskeletal Atlas II – Motor proteins and adaptors.***
Protein-level intensities for Cytoskeletal species catalogued as "motors" in Supplemental File 3 (Perseus) and Supplemental File 18.

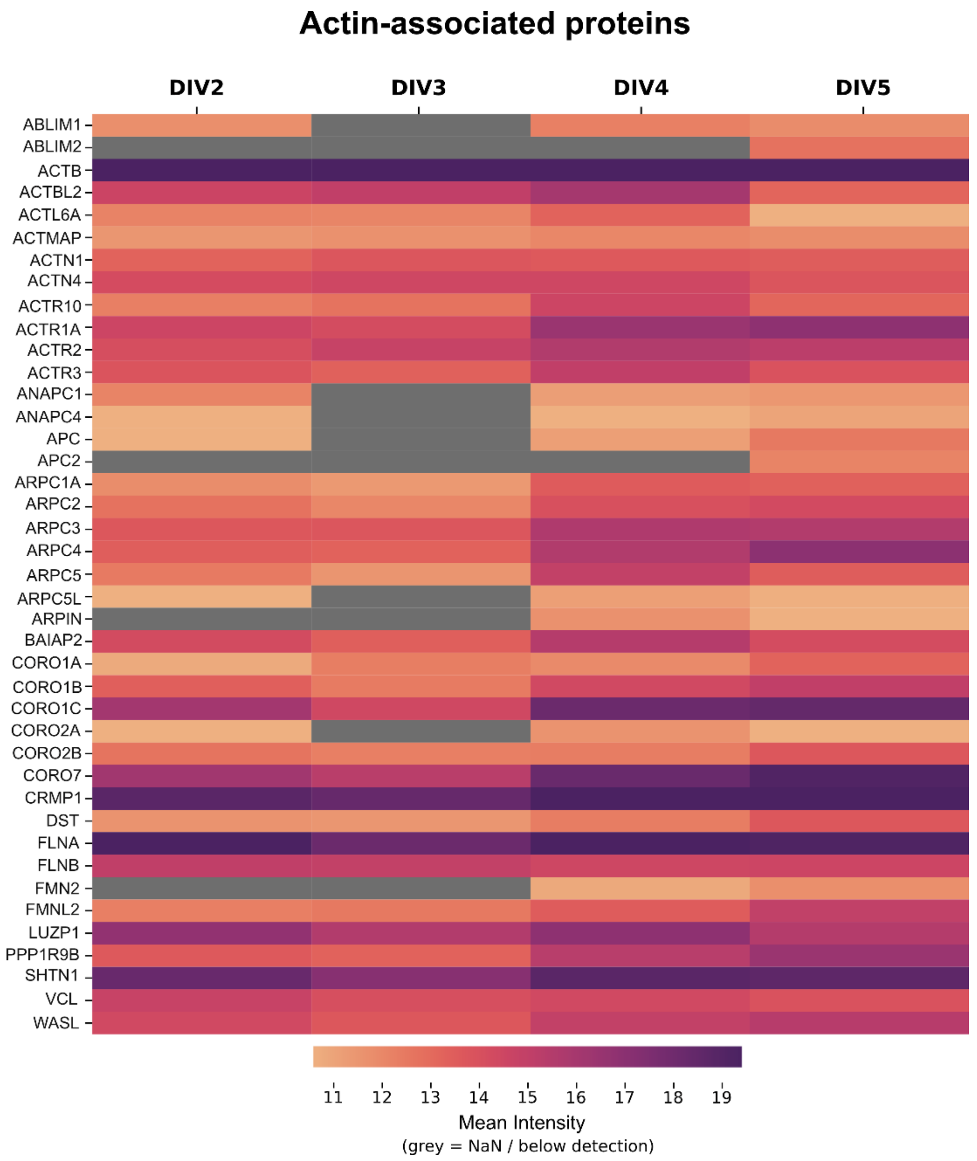


**Figure 3 – Supplement 3:** ***Cytoskeletal Atlas III – Actin-binding and associated proteins.***
Protein-level intensities for Cytoskeletal species catalogued as "actin" in Supplemental File 3 (Perseus) and Supplemental File 18.

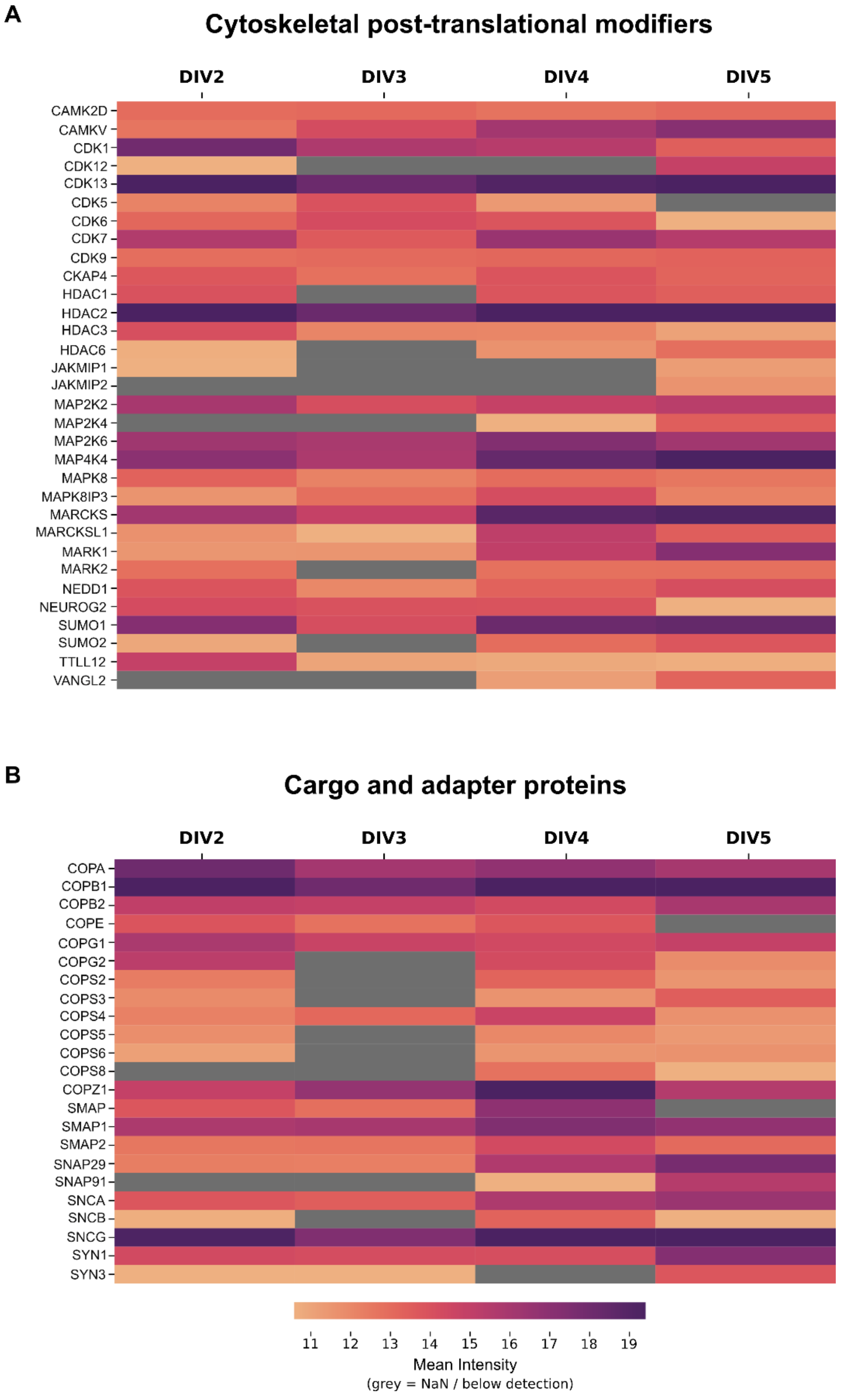


**Figure 3 – Supplement 4:** ***Cytoskeletal Atlas IV – PTM (A) and cargo-associated (B) proteins.***
Protein-level intensities for Cytoskeletal species catalogued as "regulatory" (A) and "cargo" (B) in Supplemental File 3 (Perseus) and Supplemental File 18.

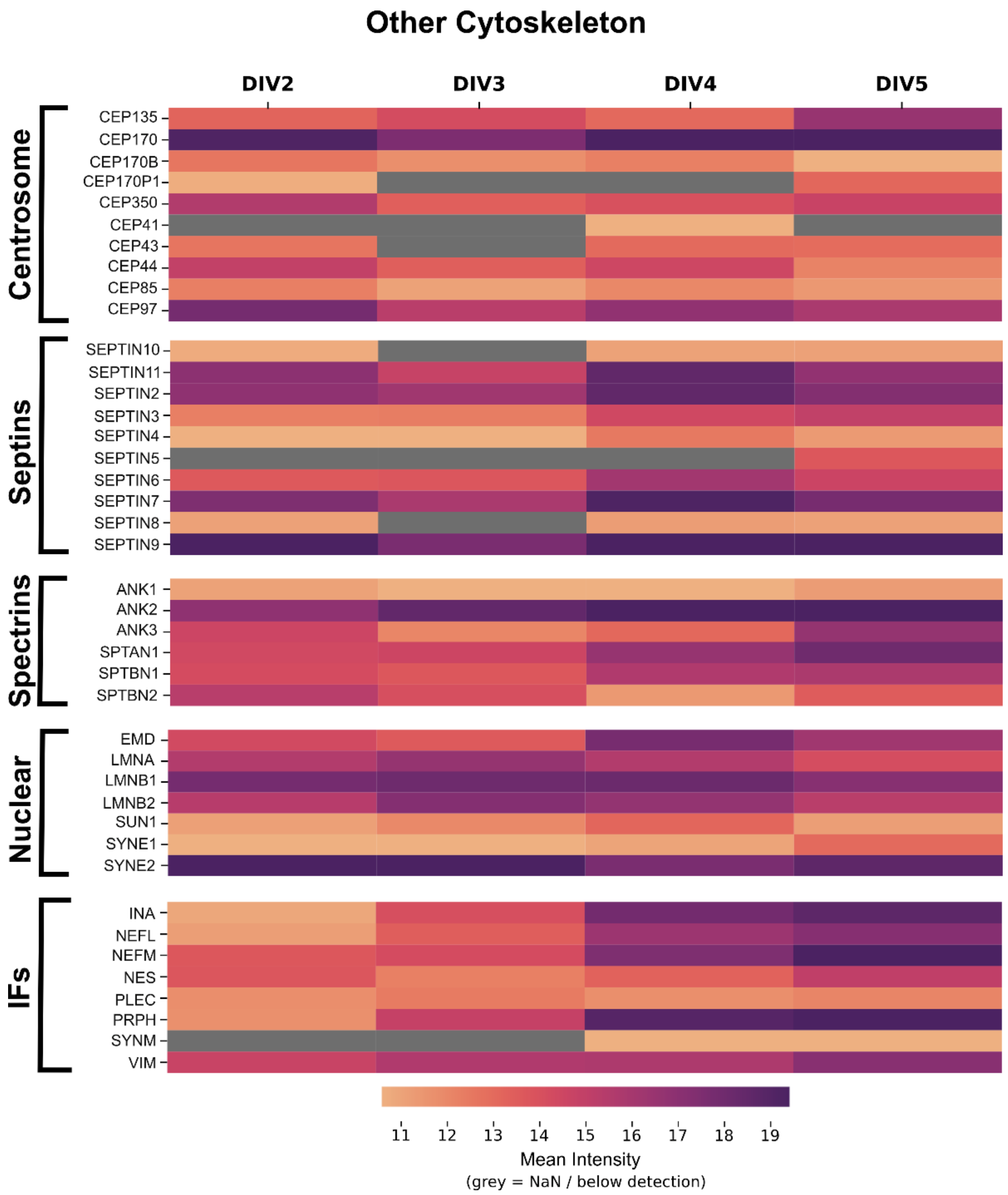


**Figure 3 – Supplement 5:** ***Cytoskeletal Atlas V – Associated cytoskeletal systems.***
Protein-level intensities for Cytoskeletal species catalogued as “IF”, “Nuclear”, “Spectrins”, “Septins”, or “Centrosome” in Supplemental File 3 (Perseus) and Supplemental File 18.

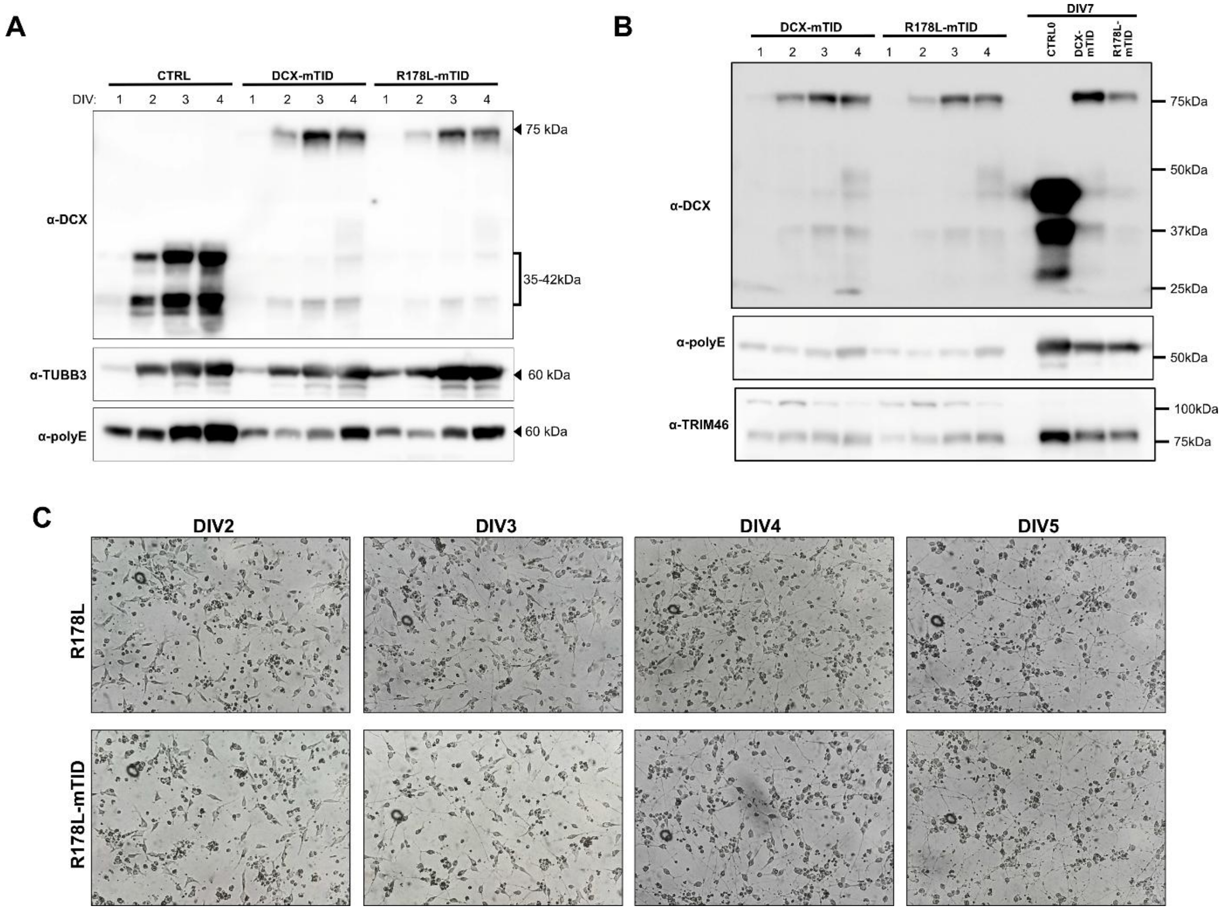


**Figure 4 – Supplement 1:** ***R178L-mTID knock-in and cell line validation.*** **(A)** Immunoblot showing expression of DCX and TUBB3, and the level of polyglutamylated tubulin across DIVs in CTRL0, DCX-mTID and R178L-mTID cells. **(B)** Immunoblot demonstrating the levels of tubulin polyglutamylation and TRIM46 expression relative to DCX expression in each cell line at DIV7. **(C)** Differentiation series of polarized light microscopy images comparing morphology of R178L mutant cells with knock-in R178L-mTID across DIVs.

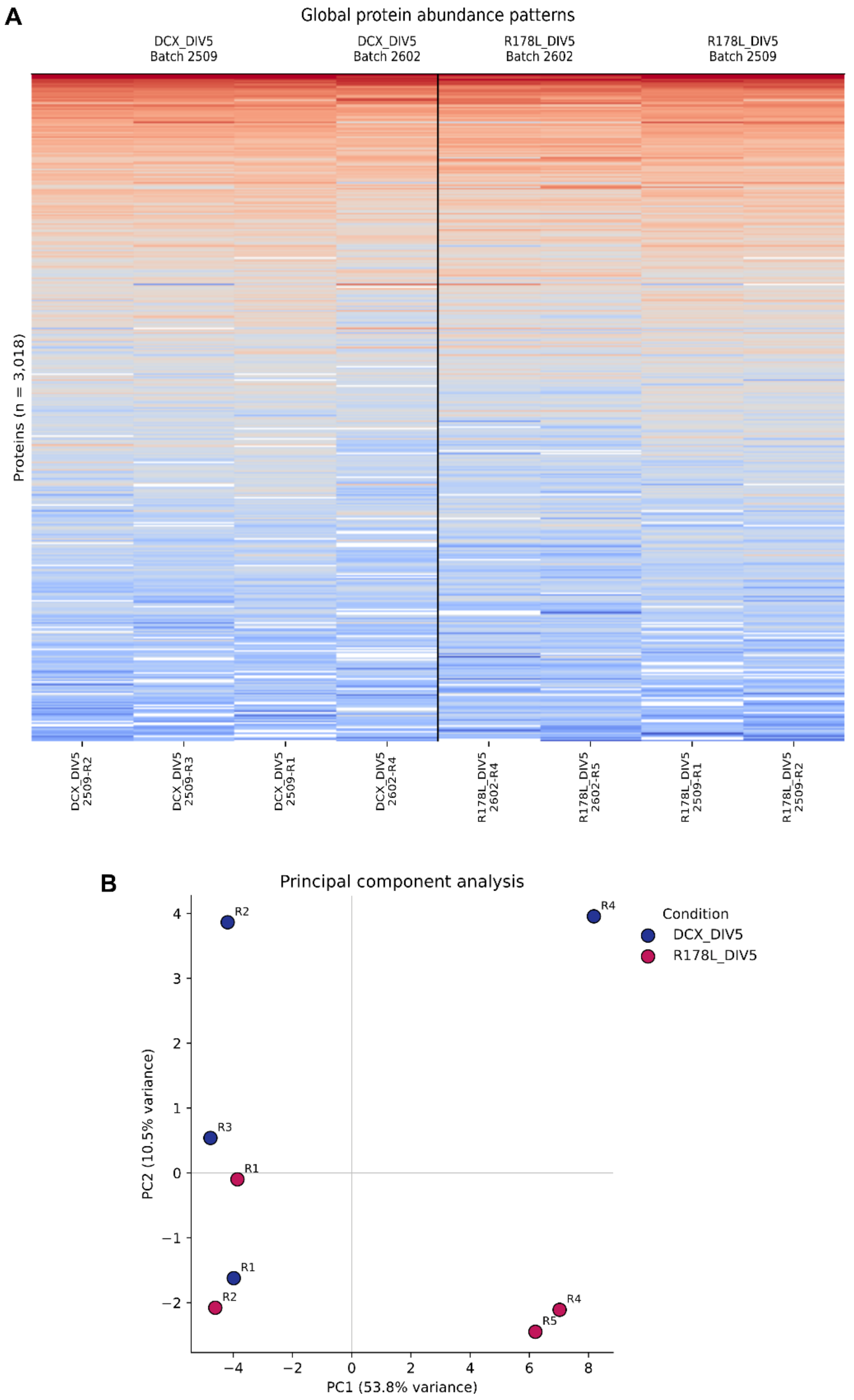


**Figure 4 – Supplement 2:** ***Batch-level and replicate behavior of DIV5 LC-MS samples*****. (A)** Heat map showing hierarchical clustering of protein intensities across individual DIV5 WT and R178L samples. Biological replicates (R1–R5; R3 excluded for poor data quality) and MS acquisition batches (2509 and 2602) are indicated for DIV5 WT-DCX (n = 4), and DIV5 R178L-DCX (n = 5). **(B)** Principal component analysis (PCA) showing clustering of biological replicates between conditions and batches.

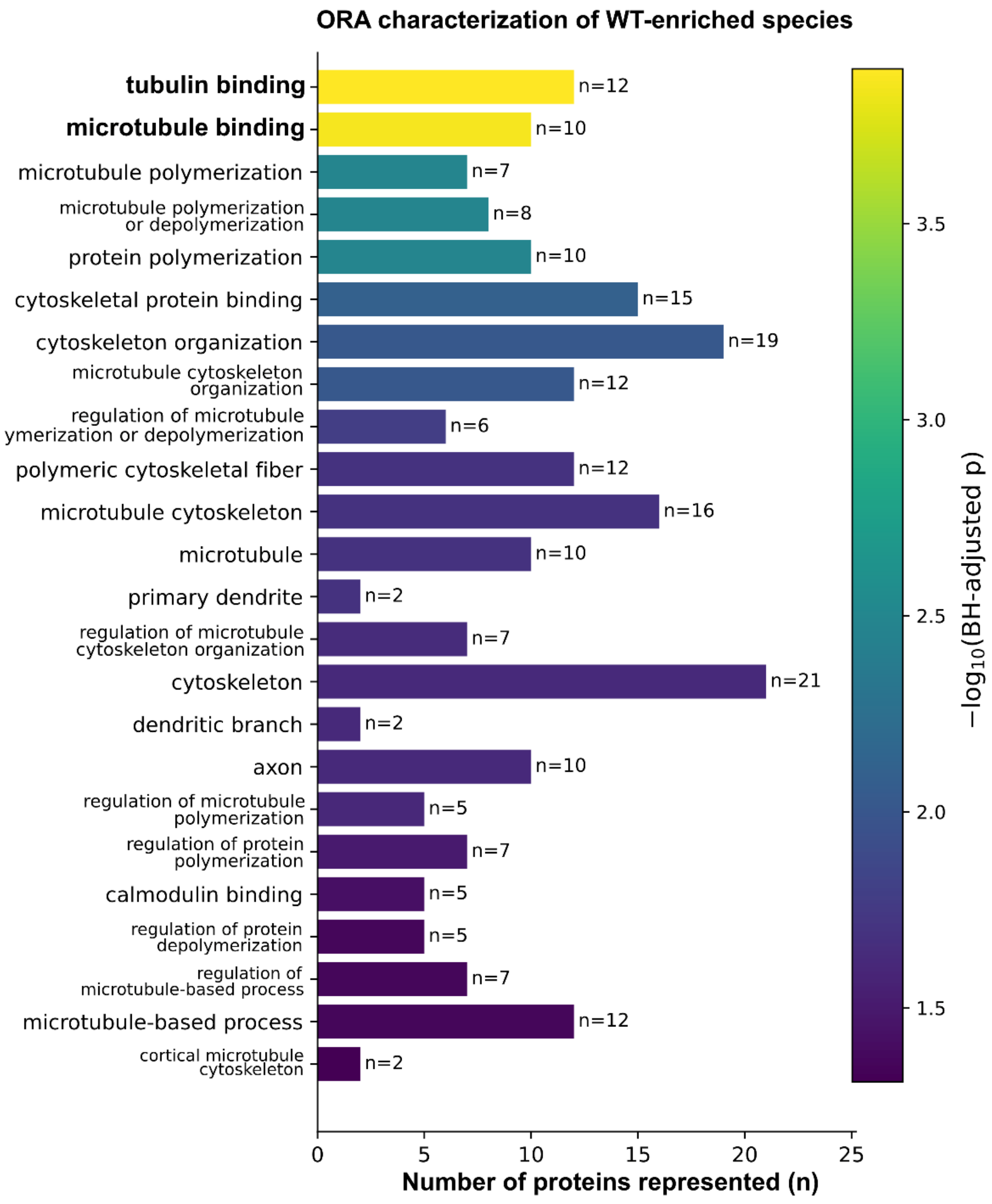


**Figure 5 – Supplement 1:** ***Gene Ontology over-representation analysis (GO-ORA) of WT-enriched proteins.*** GO enrichment plot showing over-representation of proteins involved in microtubule-binding and dynamics regulation. Bar length represents the number of proteins from the curated cytoskeletal subset annotated to each GO term (n); Bar colour represents enrichment significance, expressed as $-\log_{10}$ of the Benjamini-Hochberg-adjusted *p*-value. Input list is represented by all proteins enriched >1-fold ($\log_2$FC > 0) relative to DCX(R178L). Full list of terms is supplied in **Supplemental File 22**.

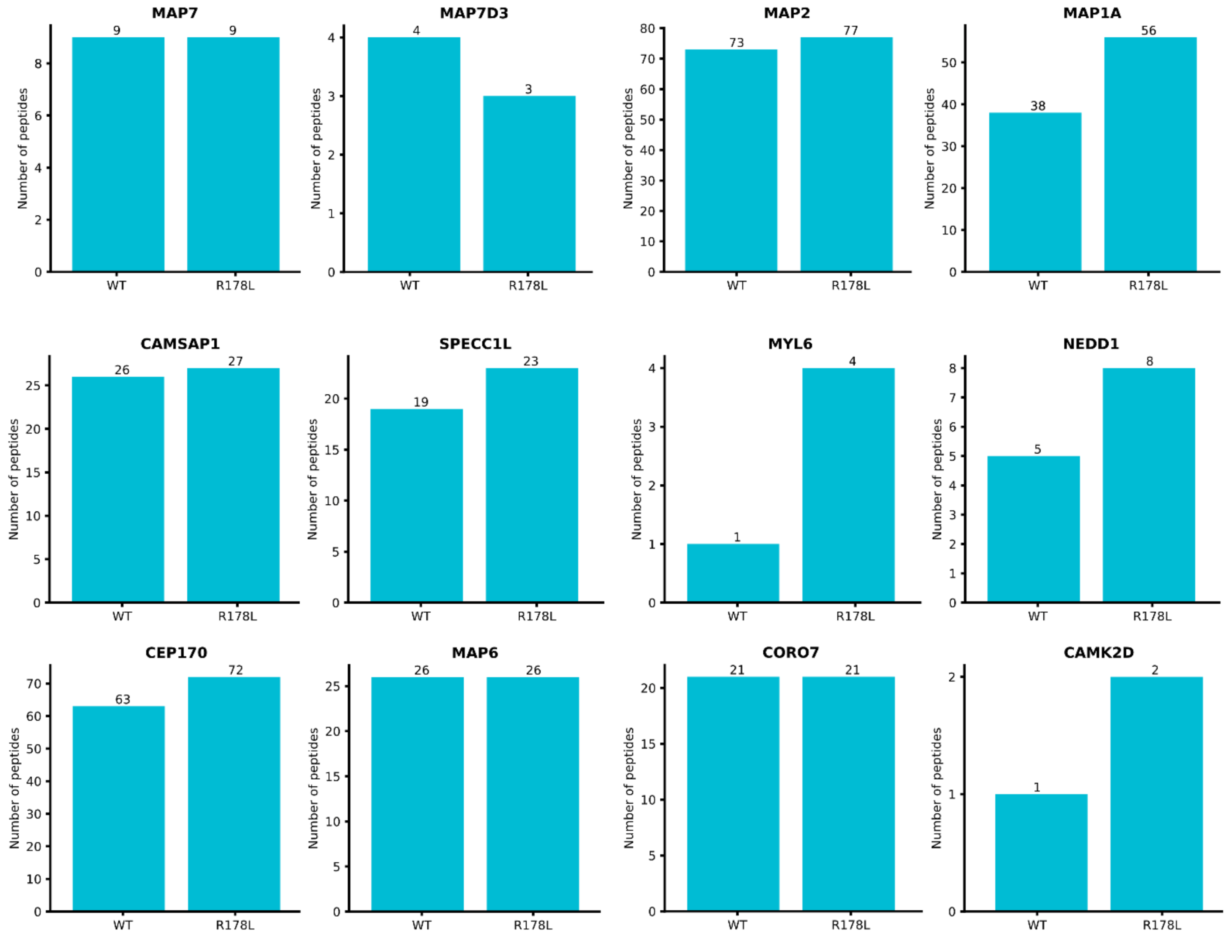


**Figure 5-Supplement 2:** ***Peptide detection of WT-enriched cytoskeletal proteins.***
Bar charts counting number of peptides detected in DCX(WT) vs DCX(R178L) condition, for Cytoskeletal proteins which are found to be significantly enriched in the WT condition at the protein level. Peptide counts are summarized atop each bar. Peptides with >3 valid values across condition replicates were not counted.

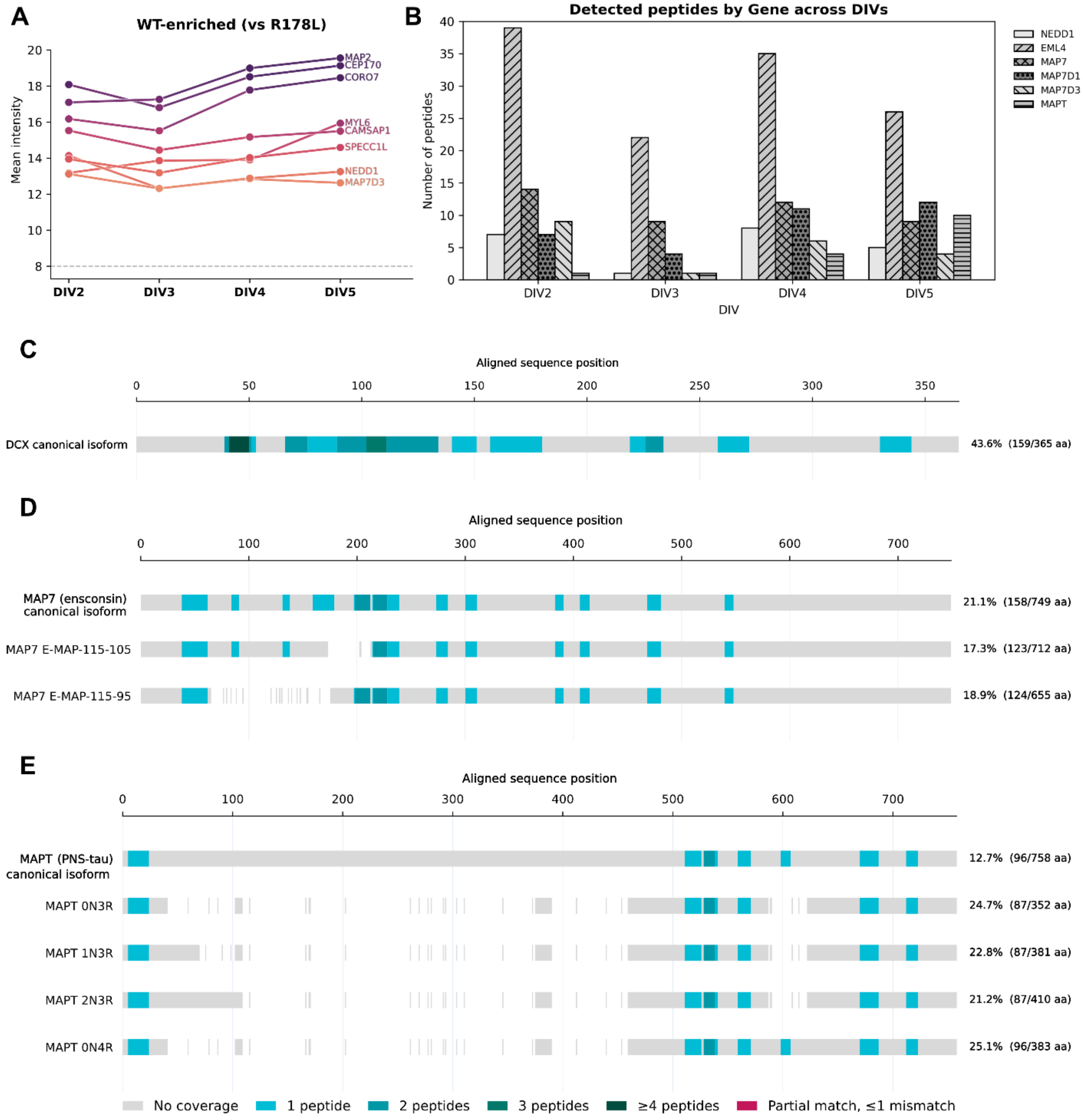


**Figure 5-Supplement 3:** ***Peptide detection across DIVs and R178L.*** **(A)** Line plot showing protein-level intensity of WT-enriched proteins across DIV2-5. **(B)** Bar chart showing the number of unique peptides detected for listed proteins, across DIV2-5. **(C-E)** Sequence alignments with LC-MS peptides detected for DCX (B), MAP7 (C), and MAPT (D) showing overlap between canonical and alternative isoforms.

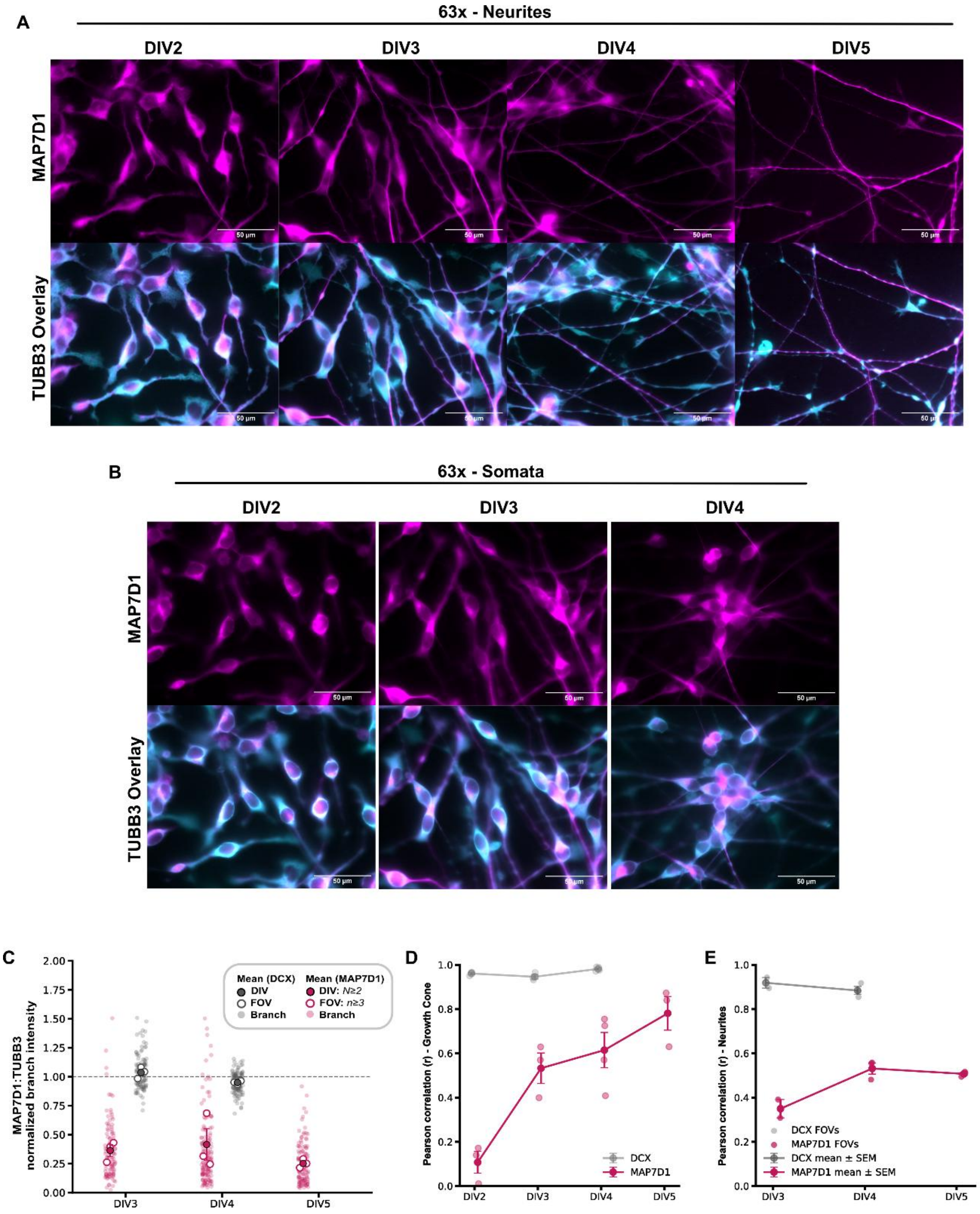


**Figure 6-Supplement 1:** ***MAP7D1 spatial relationship with TUBB3 and DCX throughout differentiation.*** **(A-B)** Widefield microscopy images acquired at 63x with showing overlay of MAP7D1 (magenta) and TUBB3 (cyan) in neurite processes (A) and in somata (B) at DIV2-5. **(C)** Swarm plot illustrating signal intensity ratios of MAP7D1 (magenta) and DCX (grey) to TUBB3 for individual branch sites ("Branch"), the mean per field ("FOV"), and the mean per DIV ("DIV"), measured across 2 biological replicates, and >3 representative fields. Scale bars are 50 µm. **(D)** Pearson coefficients per field calculated for signal overlap in growth cone ROIs; swarmplot shown in **Figure 7E**. **(E)** Pearson coefficients per field calculated for signal overlap along neurites; line profiles shown in **Figure 7C.**

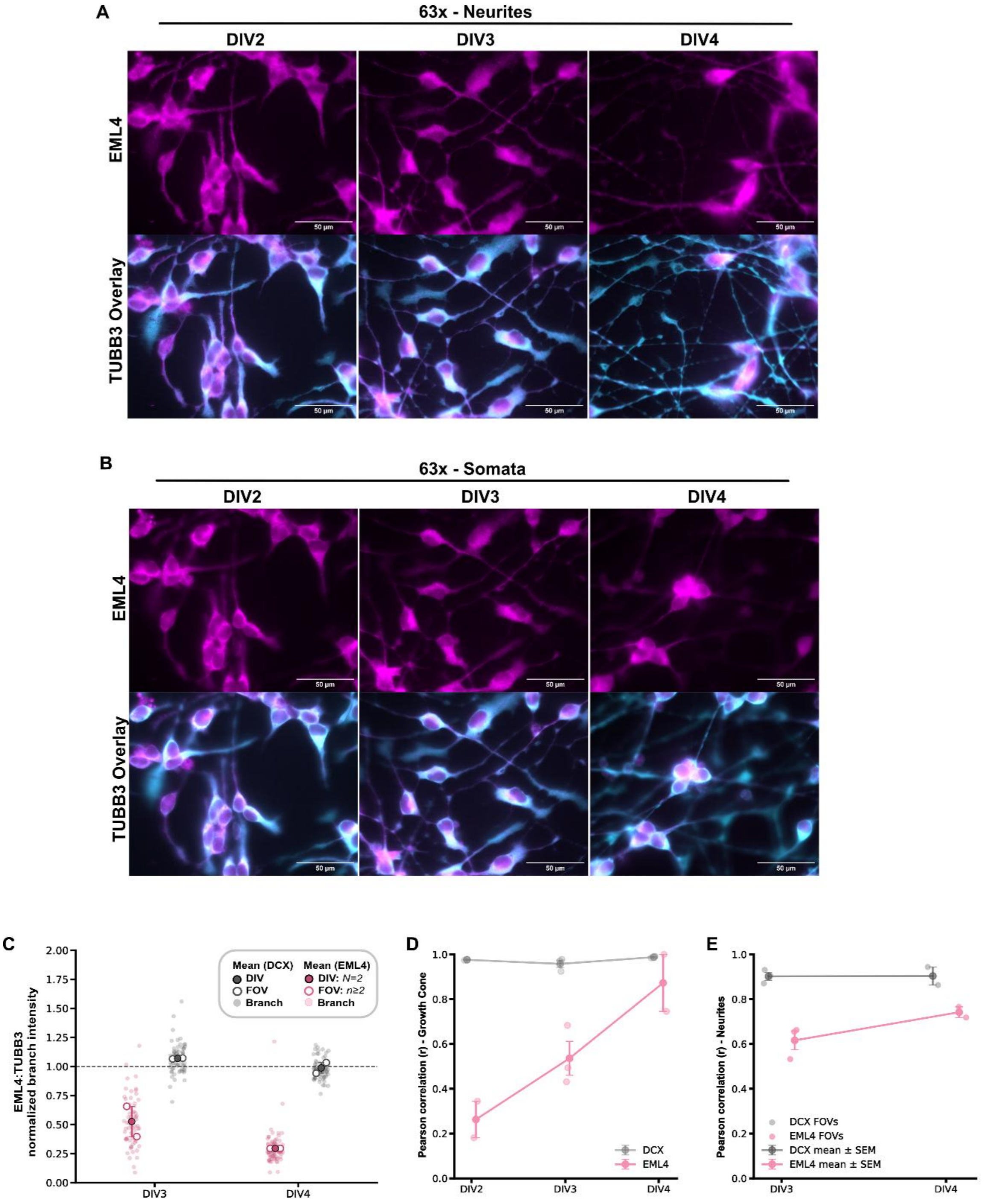


**Figure 6-Supplement 2:** ***EML4 spatial relationship with TUBB3 and DCX throughout differentiation.*** **(A-B)** Widefield microscopy images acquired at 63x with showing overlay of EML4 (pink) and TUBB3 (cyan) in neurite processes (A) and in somata (B) at DIV2-4. **(C)** Swarm plot illustrating signal intensity ratios of EML4 (pink) and DCX (grey) to TUBB3 for individual branch sites (“Branch”), the mean per field (“FOV”), and the mean per DIV (“DIV”), measured across 2 biological replicates, and >3 representative fields. Scale bars are 50 µm. **(D)** Pearson coefficients per field calculated for signal overlap in growth cone ROIs; swarmplot shown in **Figure 7F**. **(E)** Pearson coefficients per field calculated for signal overlap along neurites; line profiles shown in **Figure 7D.**
